# Verification of our empirical understanding of the physiology and ecology of two contrasting plantation species using a trait database

**DOI:** 10.1101/2021.07.23.453564

**Authors:** Yoko Osone, Shoji Hashimoto, Tanaka Kenzo

**Author notes:** Correspondence to: Tanaka Kenzo.

## Abstract

The effects of climate change on forest ecosystems take on increasing importance more than ever. Information on plant traits is a powerful predictor of ecosystem dynamics and functioning. We reviewed the major ecological traits, such as foliar gas exchange and nutrients, xylem morphology and drought tolerance, of *Cryptomeria japonica* and *Chamaecyparis obtusa*, which are major timber species in East Asia, especially in Japan, by using a recently developed functional trait database for both species (SugiHinokiDB). Empirically, *C. obtusa* has been planted under drier conditions, whereas *C. japonica* has been planted under wetter conditions. Our analyses revealed followings: The maximum photosynthetic rate, stomatal conductance, foliar nutrient content and soil-to-foliage hydraulic conductance were higher in *C. japonica* than in *C. obtusa* and were consistent with the higher growth rate of *C. japonica*. In contrast, the foliar turgor loss point and xylem pressure corresponding to 50% conductivity, which indicate drought tolerance, were lower in *C. obtusa* than in *C. japonica* and are consistent with the drier habitat of *C. obtusa*. Ontogenetic shifts were also observed; as the age and height of the trees increased, many foliar nutrient concentrations decreased, and the foliar minimum midday water potential and specific leaf area also decreased. This suggests that an ontogenetic reduction in photosynthesis occurred due to an increase in drought stress with tree height and age. However, among the Cupressaceae worldwide, the drought tolerance of *C. japonica* and *C. obtusa* is not as high. This may be related to the fact that the Japanese archipelago has historically not been subjected to strong dryness. The maximum photosynthetic rate showed intermediate values within the family, indicating that *C. japonica* and *C. obtusa* exhibit relatively high growth rates in the Cupressaceae family, and this is thought to be the reason why they have been selected as economically suitable timber species in Japanese forestry. This study clearly demonstrated that the plant trait database provides us a promising opportunity to verify out empirical knowledge of plantation management and helps us to understand effect of climate change on plantation forests by using trait-based modelling.

## 1 Introduction

There is an emerging scientific consensus that the global climate change is resulting in decreased stability in forest ecosystems (Mann et al., 1998; Allen et al., 2010). The effects of climate change on the forestry sector have been examined in general terms for many regions of the world but rarely with sufficient temporal or spatial resolution to influence regional or local forest management (IPCC, 1996; Roos, 1996; Sykes and Prentice, 1996; Bradshaw et al., 2000). The major issues include how the ranges in which commercially important tree species are suitable for plantations will change in the future and whether climatic influence can be overridden by appropriate forest management. Answering these questions requires a basic understanding of the physiology and ecology of target tree species.

Information on plant traits, that is, any physiological, morphological or phenological features measurable at the individual level (Violle et al., 2007), is now widely used to predict how forests will respond to future climate change (White et al., 2002; Sato et al., 2007; Zaehle et al., 2010; Kattge et al., 2020). Process-based models generally use leaf-scale process, such as, photosynthetic capacity, stomatal response to vapor pressure deficit (VPD) and respiration of a plant species or a functional type for modelling C dynamics under given climate scenarios (Chapin et al., 2011; Toriyama et al., 2021). There are also studies focusing on traits more directly related to drought sensitivity for predicting future hydraulic risk. For example, leaf water potential at turgor loss (Ψtlp), which had been recognized a classical index of plant water stress, was demonstrated to be a powerful indicator of drought tolerance within and across biomes (Bartlett et al., 2012; Peters et al., 2021), while hydraulic safety margins, defined as difference between minimum xylem water potential and water potential at which 50% loss of conductivity occurs (Ψ50) is becoming widely used for a predictor of drought-induced tree mortality (Choat et al., 2012; Anderegg et al.; 2016). Clearly, trait information holds promise for better understanding of the vulnerability to drought, as well as parameterizing models with increased robustness and accuracy.

With the increasing demands for trait information, trait databases are becoming key research tool in this study field (Kattge et al., 2020). One of the strengths of trait databases is that they provide a wide array of traits for a species all at once, which is generally difficult in a single study since measurements of physiological and morphological properties are time- and labour-intensive. Another advantage is that they show the variability within a species since they stores data from different studies which measured plants at different ages in different locations. Many functional traits change ontogenetically as plants grow (Beets and Madgwick, 1988; Bargali et al., 1992; Bond, 2000; Niinemets, 2002; Hooker and Compton, 2003; Polglase et al., 2006; Yang and Luo 2011). Understanding the ontogenetic drift of key functional traits is important for impact assessments of climate change since forest management is a long-term commitment and requires optimality of adaptation strategies at each growth stage (Niinemets, 2010; D’Amato et al., 2011).

Recently, we created a trait database for Japanese cedar (*Cryptomeria japonica* D. Don, *Cupressaceae*) and Japanese cypress (*Chamaecyparis obtusa* (Siebold et Zucc.) Endl., *Cupressaceae*) (SugiHinokiDB), which contains 24683 data for 177 plant traits compiled from diverse sources, such as papers, bulletins, reports and books (Osone et al., 2020). *C. japonica* and *C. obtusa* produce high-quality wood and have been the most important commercial tree species in Japan. They were also introduced for timber production in many regions of the world: China, Korean Peninsula, India, Nepal, Azores and Réunion (Government of Azores Islands; Rull et al., 2017; Rai and Schmerbeck, 2018). In Japan, these species were planted to forest sites according to empirically derived rule for species selection. *C. japonica*, which grows faster but thought to be less drought tolerant than *C. obtusa*, is traditionally planted on moist and nutrient-rich sites, whereas *C. obtusa* is planted on relatively dry and nutrient-poor sites (Mashimo, 1960; Hayashi, 1969; Sato, 1971). Since these management practices had worked well until recently, we have paid little attention on the physiological mechanisms underlying their habitat preferences. However, without the knowledge, we cannot predict how the species respond to climate change, nor what the optimal adaptation strategies are. SugiHinokiDB, which compiled traits that are closely related to the life history strategy (Díaz et al., 2004; Grime, 2006; Kattge et al., 2011, 2020; Kleyer et al., 2008), with a special focus on traits related to water relations, may offer a comprehensive characterization of the growth and hydraulics of these species.

In this study, using the SugiHinoki plant trait DB, we verify the empirical knowledge that *C. japonica* grows faster but is less tolerant to drought than *C. obtusa* based on three steps:

(1) We selected 20 traits that are central to the leading dimensions of plant strategy and quantified the differences in those traits between the two species. Our hypothesis is that *C. japonica* has relatively pioneer-like properties, i.e., a higher gas exchange rate, specific leaf area (m2 g-1, SLA), and xylem and foliar water conductivity, whereas *C. obtusa* shows more conservative resource use and a higher drought tolerance.
(2) We also examined the ontogenetic changes in some foliage traits. There is still limited knowledge on age or height depending changes in foliage traits, particularly their species patterns. We demonstrated how the ontogenetic patterns of key foliage traits are different between drought intolerant and drought tolerant species.
(3) Finally, we compare some hydraulic properties of these species with those of Cupressaceae worldwide. Cupressaceae species are thought to differentiate along an aridity gradient and vary greatly in their drought sensitivity. We discussed the adaptive strategies of *C. japonica* and *C. obtusa* in light of the phylogenetic lineage and potential as timber species under future climate.

## 2 Materials and Methods

### 2.1 Plant species

Japanese cedar (*Cryptomeria japonica* (L.f.) D. Don, Sugi cedar), an evergreen conifer, is the only species of the genus *Cryptomeria* in Cupressaceae. It is distributed mainly in Japan, but *C*. *fortunei*, which is genetically identical to *C. japonica* is scarcely distributed in Zhejiang, China (Tsumura et al., 1995). In Japan, its natural range is Lat. 30°to 40° with mean annual precipitation > 1800 mm (Tsukada 1982). *C. japonica*, which grows rapidly with a maximum height greater than 50 m, has been used for timber production since the prehistoric period. At present, it dominates approximately 45% of the forest area in Japan. Japanese cypress (*Chamaecyparis obtusa* (Sieb. et Zucc.) Endl.), distributed in Japan and Taiwan, is also an evergreen conifer in the *Cupressaceae* family. The northern limit of its natural range (Lat. 30°-37°) is lower in latitude than that of *C*. *japonica*. Due to low snow resistance, the species rarely appear coastal area of Sea of Japan, where there is plenty of snow in winter. Although *C*. *obtusa* grows slower than *C*. *japonica*, it produces high-quality wood and thus has also long been a commercially important species in Japan. *Chamaecyparis obtusa* dominates 15% of the forest area in Japan.

### 2.2 Plant trait database

The sugi-hinoki database (SugiHinoki DB) consists of 24683 data entries for 177 traits of *C*. *japonica* and *C. obtusa* from 364 primary sources including journal papers, unpublished data and grey literature (e.g., reports, theses, articles) (Osone et al., 2020). The traits, grouped into 15 categories by their features (Table 1), are those that are widely agreed on as relevant to plant life-history strategies, vegetation modelling and global change responses (Grime, 1977; Díaz et al., 2004; Kleyer et al., 2008; Kattge et al., 2011). Because of the limited distribution of the species, data were mainly obtained from forest sites in Japan (30°20’N, 130°32’E - 41°25’N, 140°6’E) but also from arboretums or plantations in Taiwan, Korea and China. The database includes data from plants grown in plantation forests, natural forests, and those grown under experimental conditions. Each data entry is accompanied by ancillary information about the location, environmental conditions, experimental treatment, measurement methods, status of measured individuals in the stand and the position of measured parts (e.g., upper or lower crown for photosynthetic measurements). Further details on the database are given in Osone et al. (2020).

**Table 1.**
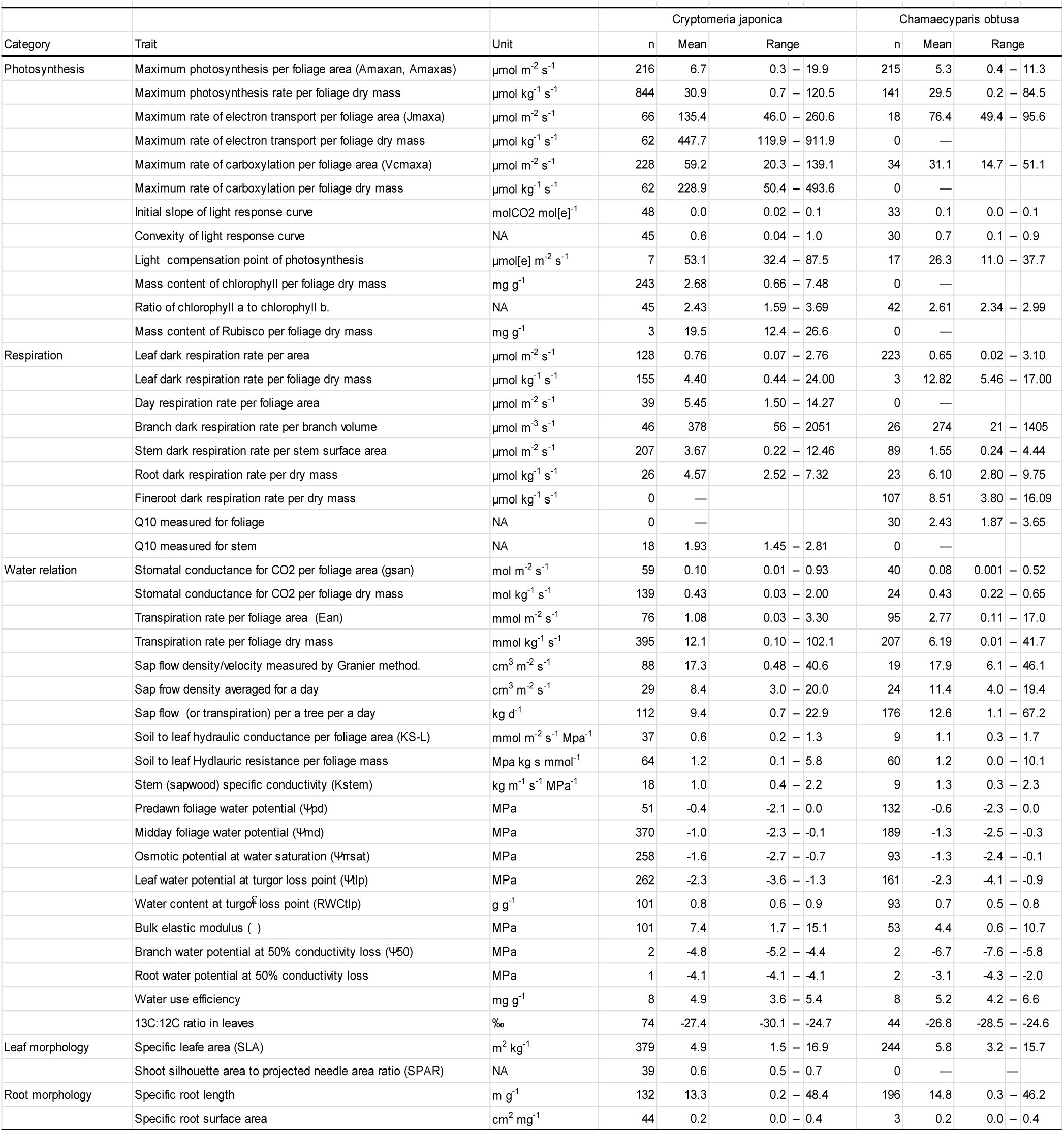
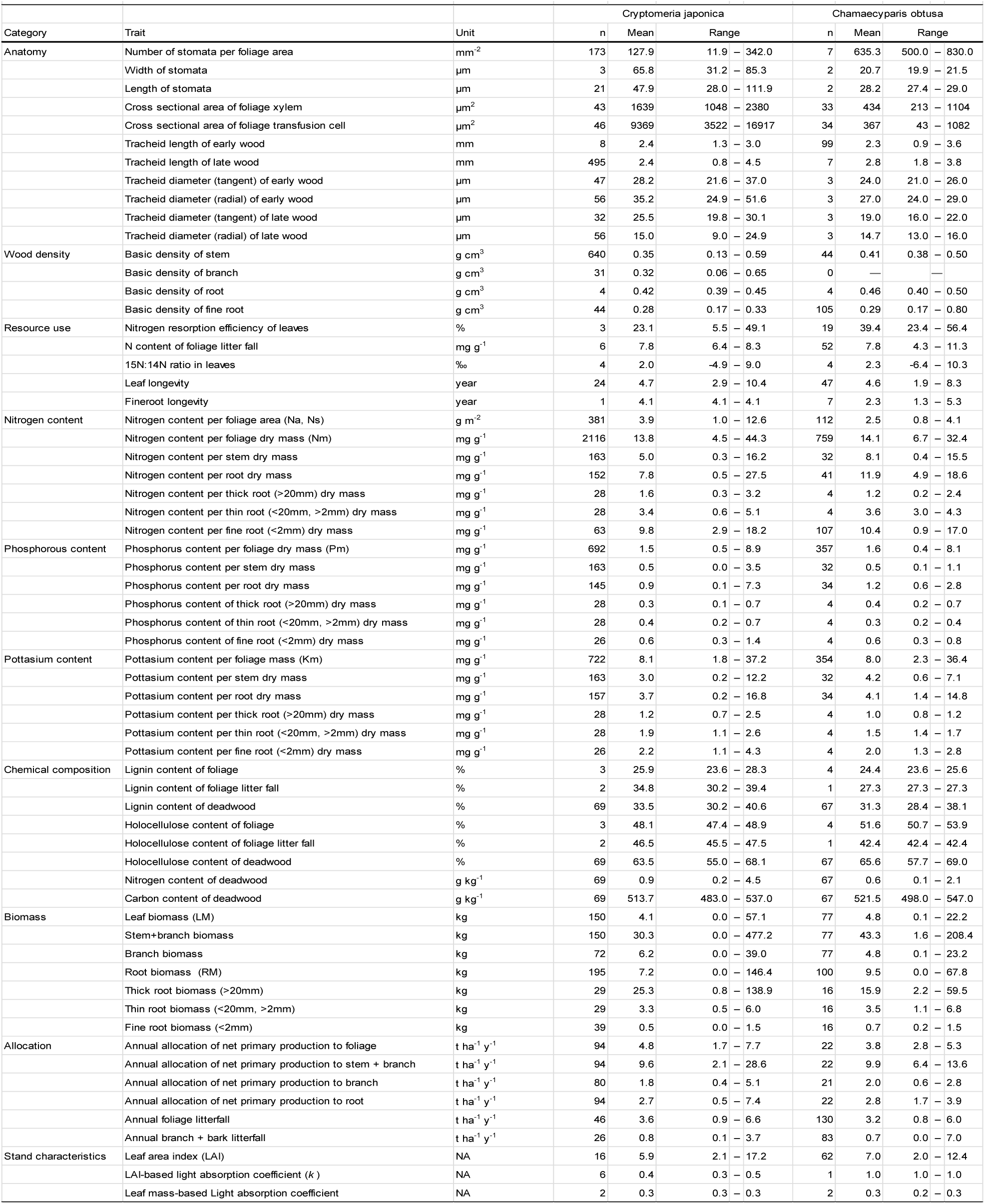
List of 108 major plant traits selected from the SugiHinoki DB. Number of data entries and means and ranges were calculated before outliers were removed for subsequent analyses.

Table 1 lists 108 major plant traits selected from the SugiHinoki DB (Osone et al., 2020). The traits with the highest number of data entries are mass-based nutrient contents such as mass-based foliar N concentration (N_m,_ 2875), K concentration (K_m_, 1076) and P concentration (P_m_, 1049), which had once been easy-to-measure indices of plant physiological status. Other basic foliar properties, including photosynthetic capacities (Amax_a_, 431; Amax_m_, 985), SLA (623), foliar pressure-volume curve parameters (Ψ_tlp_, 423; RWC_tlp_ 194; Ψ_πsat_, 351; ℇ, 154) and midday (minimum) foliage water potential (Ψ_md_, 559), are also those with the highest number of data entries. On the other hand, fewer data are available for properties that require more complex measurements, such as water conductance/conductivity and the hydraulic safety during water transport. *C*. *japonica* and *C*. *obtusa* showed similar patterns in data abundance among traits, but *C*. *japonica* had more data for most traits (81 out of 108 traits). Each trait of a species showed quite large variation in the values since the database contains data for trees of different ages, grown under different conditions and measured at different times of the day or year. The data distribution of each trait showed a convex curve when data were abundant, but distributions were mostly positively skewed.

### 2.3 Selection of traits for the comparison of *C. japonica* and *C. obtusa*

To detect the ecophysiological basis for the empirical knowledge that *C. japonica* grows faster on nutrient-rich moist soil than *C. obtusa*, we selected 20 traits that are considered relevant to plant growth and water relations from SugiHinokiDB. Those include the maximum photosynthetic rate per area (Amax_a_), maximum carboxylation rate per area (Vcmax_a_), maximum electron transport rate per area (Jmax_a_), foliar dark respiration rate per area (R_a_), stomatal conductance for CO_2_ per area (gs_a_), foliar N per area (N_a_), SLA, foliar water potential at the turgor loss point (Ψ_tlp_), foliar relative water content at the turgor loss point (RWC_tlp_), foliar osmotic potential at full turgor (Ψ_πsat_), bulk elastic modulus (ε), soil-to-foliage water conductance (K_S-L_), stem specific conductivity (K_stem_), tracheid diameter of the stem, tracheid length of the stem, basic density of the stem, xylem water potential at a 50% loss of conductivity (Ψ_50_), foliage mass (LM), stem mass (SM) and root mass (RM). Abbreviations and units of the traits used in the analyses are shown in Table 1.

### 2.4 Selection of traits for the analysis of foliage age or height dependency

For the analysis of age (size) dependency, we used the foliage nitrogen content per foliage dry mass (N_m)_, foliage phosphorus content per foliage dry mass (P_m_), foliage potassium content per foliage dry mass (K_m_), specific leaf area (SLA) and midday foliage water potential (Ψ_md_). These are key traits for plant growth and water relations, and changes in these traits with age or size could have considerable effects on stand growth, carbon and nutrient cycling, and thus forest management. In SugiHinoki DB, they are abundant in data entries from many sources where measurements were performed for many trees of different ages or sizes. However, since the data were not obtained from a single carefully controlled experiment but were the compilation of multiple studies, we could not separate size and age effects that may independently affect foliage traits. Thus, we assessed the dependency by the strength of the correlation between each trait and age and height.

### 2.5 Two measures of foliage-area based traits

Shoots of *C*. *japonica* have complex structures, with needles being attached densely and helicoidally to a stalk, while shoots of *C*. *obtusa* are planar with scale-like leaves arranged on a flat surface (Fig. 1). As a result, in *C*. *japonica*, there are large differences in projected needle area and the shoot silhouette area, and thus the area-based foliar traits differ largely depending on whether the ‘foliage area’ is projected needle (or scale) area (A_n_) or the shoot silhouette area (A_s_). The relationship between the trait values presented on a needle area basis (T_leaf_/A_n_) and shoot silhouette area basis (T_leaf_/A_s_) for any trait (T_leaf_) is given as

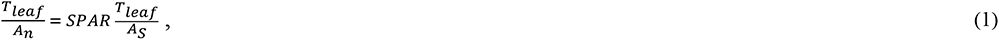

**Fig. 1.**
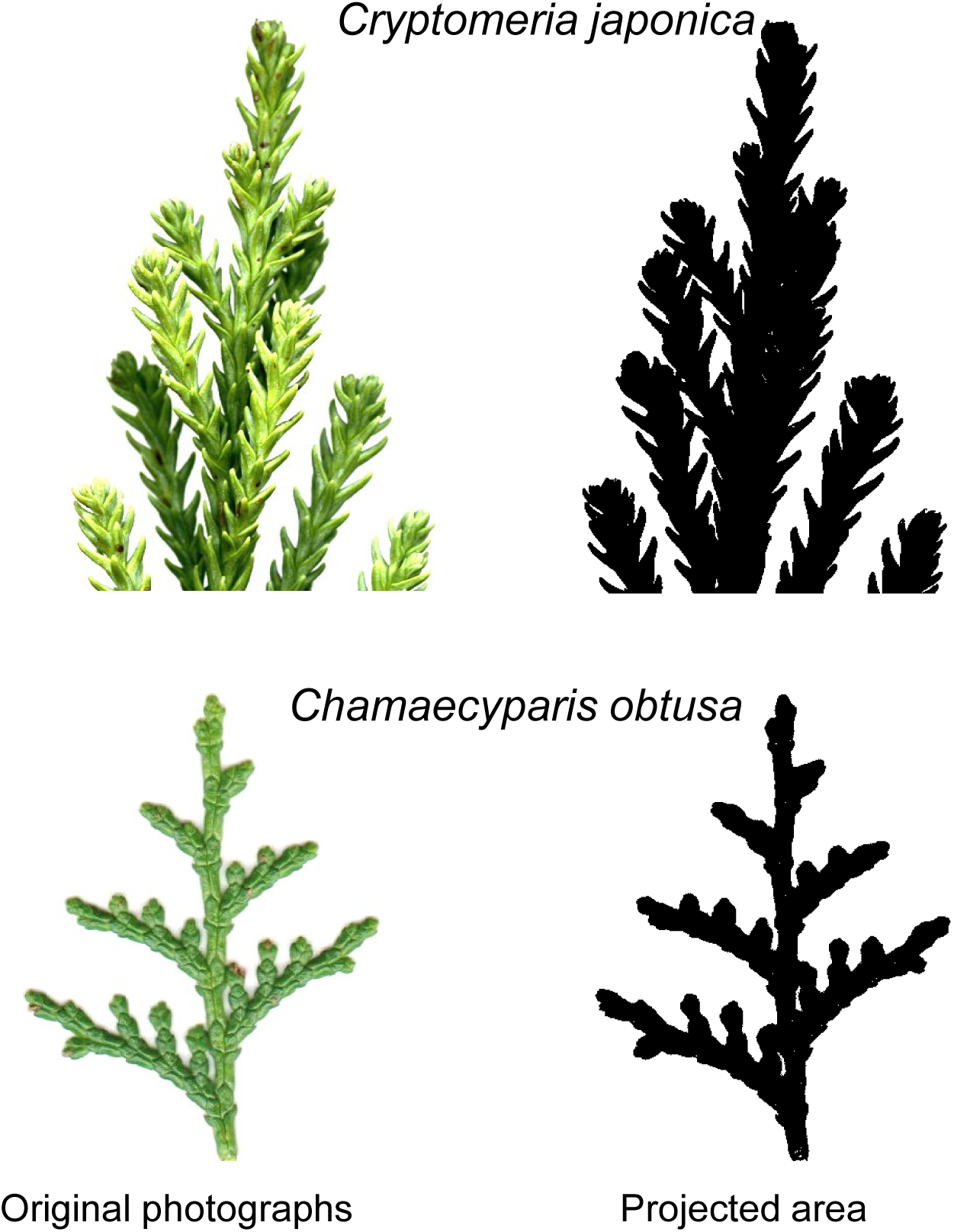
Photographs of *Cryptomeria japonica* (top) and *Chamaecyparis obtusa* (bottom). The right side is an example of the foliar projected area. Note that the projected area is underestimated in *C*. *japonica* because the foliage of *C*. *japonica* has a complex three-dimensional structure.

where SPAR is the shoot silhouette and projected needle area ratio (Stenberg, 1996; Ishii et al., 2007) and is given as

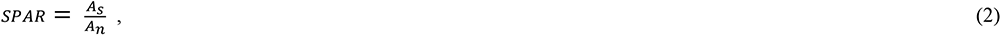

The SPAR is less than 1 in conifers with complex shoot structures and is 1 in broad-leaf species and conifers with planar shoots, such as *C. obtusa*. A smaller SPAR means higher needle clumping or higher mutual shading in a shoot. SPAR can vary among species and the crown position of a tree (Carter and Smith, 1985; Ishii et al. 2007; Thérézien et al., 2007). However, Inoue et al. (2020) found that the relationships between needle area and shoot projected area were almost consistent among shoots of different ages and different sizes among 4 cultivars of *C*. *japonica,* with a slope (=SPAR) of 0.63.

Similarly, the relationship between needle area-based SLA (An/M_n_) and shoot silhouette area-based SLA (As/M_s_) is

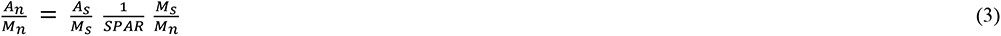

where Mn and Ms are the needle mass of a shoot and the mass of a shoot, respectively. In the database, these two values were stored individually. In this analysis, we focused on both needle/scale area-based (presented with suffix ‘n’ after trait abbreviation such as ‘Amax_an_’) and shoot silhouette area-based traits (presented with suffix ‘s’ after trait abbreviation such as ‘Amax_as_’) since the two could have different ecological meanings, especially in terms of light capture (Carter and Smith, 1985; Leverenz and Hinckley, 1990; Stenberg et al., 1995; Stenberg, 1996; Thérézien et al., 2007).

### 2.6 Data treatment for analysis

For all selected data, errors were checked, and outliers, which we defined as data out of accepted ranges, were excluded from the database. Data collected under experimental conditions (plants grown in a greenhouse, growth chamber and lysimeter; plants grown in planters; plantation forests with experimental fertilization, etc.) were also excluded, which resulted in the exclusion of most young plants (0-2 years). Gas exchange rates and traits related to water relations were filtered for the measurements conducted during summer and early autumn (Jun-Oct), except where the focus was monthly or seasonal changes.

### 2.7 Statistical analysis

The age and height dependency of the foliar traits were assessed by regression analysis. Differences in means between species were tested by Student’s t-test. All statistical analyses were performed with R Version 3.4.4 (R Core Team, 2018).

## 3 Results and Discussion

### 3.2 Differences in traits related to growth rate

#### 3.2.1 Photosynthesis

In contrast to our hypothesis that photosynthetic capacity and foliar (needle or scale) N concentration are higher in fast-growing *C. japonica* than in slow-growing *C. obtusa*, they were not significantly different between the species when presented on a foliage area basis (Amax_an_, Vcmax_an_, Jmax_an_, N_an_) (Fig. 2a-2d). However, since the two species have completely different shoot morphologies (Fig. 1), a simple comparison of these foliage area-based measurements may fail to characterize their photosynthetic properties (Smith et al., 1991; Smith and Brewer, 2002). Photosynthetic capacity is usually presented per unit foliage area on the assumption that the foliage area represents the amount of solar radiation intercepted. However, the interception of solar radiation is not directly related to the total foliage area unless foliage is oriented horizontally. In *C*. *japonica*, where needles are attached helicoidally to a stalk, mutual shading could occur within the shoots, which reduces the photosynthetic rate per needle area (Stenberg et al., 1995; Stenberg, 1996; Thérézien et al., 2007). In such shoots, as in many other coniferous shoots, light interception is determined by the shoot silhouette area rather than the total needle area, and photosynthetic characteristics should be evaluated based on both photosynthesis per needle area and per shoot silhouette area (Stenberg et al., 1995).

**Fig. 2.**
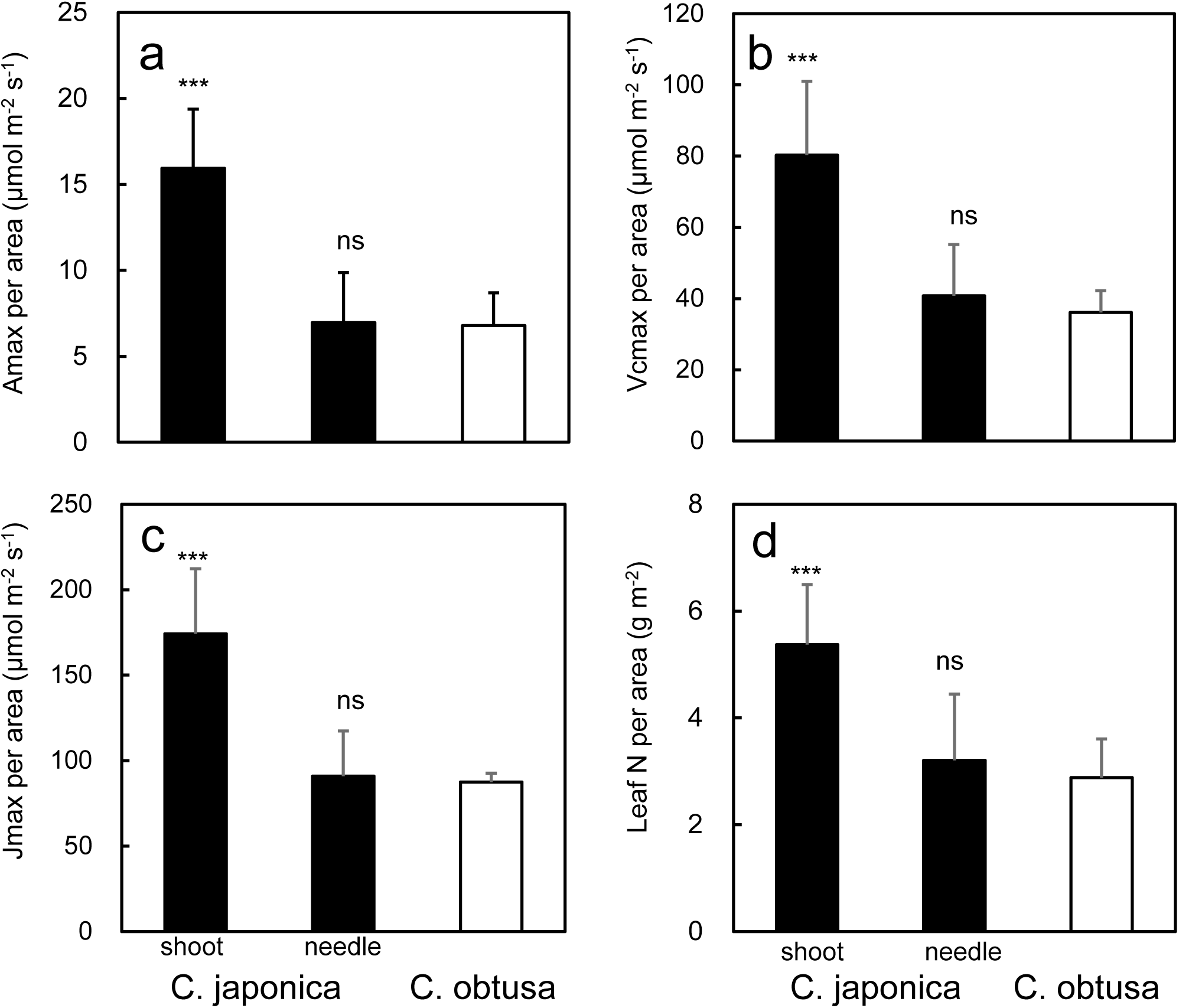
Photosynthetic properties of *C. japonica* and *C*. *obtusa*. In *C. japonica*, each trait is presented on a shoot silhouette area basis and projected needle area basis. Differences between each *C*. *japonica* trait (shoot or needle basis) and each *C*. *obtusa* trait were determined by a t-test; ***, P<0.001; xx, P<0.01; x, P<0.05; ns, not significant.

On a shoot silhouette basis, the photosynthetic capacity and foliar N contents were higher for *C. japonica* (Amax_as_, Vcmax_as_, Jmax_as_, N_as_) than for *C. obtusa* (Amax_an_, Vcmax_an_, Jmax_an_, N_an_), by 1.9 – 2.1 times (Fig. 3). The marked increases in the photosynthetic capacity of *C. japonica* when presented based on the shoot silhouette area are theoretically justified as follows. Photosynthesis per silhouette area is photosynthesis per needle area divided by SPAR (eqn. 1), which is 0.53-0.73 in *C. japonica* (SugiHinoki DB). The Amax_as_ calculated by eqn. 1 with this SPAR and Amax_an_ (7.55 μmol m^-2^ s^-1^) is 10.34-14.25 μmol m^-2^ s^-1^, which is similar to the measured Amax_as_ (Fig. 3a). Note that the two measures of photosynthesis are consistent in *C*. *obtusa* in which shoot is planar and SPAR is one. The higher Amax_as_ in *C. japonica* than in *C. obtusa* despite the similar Amax_an_ suggests that under saturating irradiance, densely packed needles on the shoots of *C. japonica* can absorb more irradiance than the planar shoots of *C*. *obtusa* and thereby achieve a higher photosynthetic rate per shoot. However, this could be at the expense of the photosynthetic efficiency of each needle in *C*. *japonica*; that is, at low irradiance, the photosynthetic rate per needle area could be decreased more than that of *C*. *obtusa* due to mutual shading of needles.

**Fig. 3.**
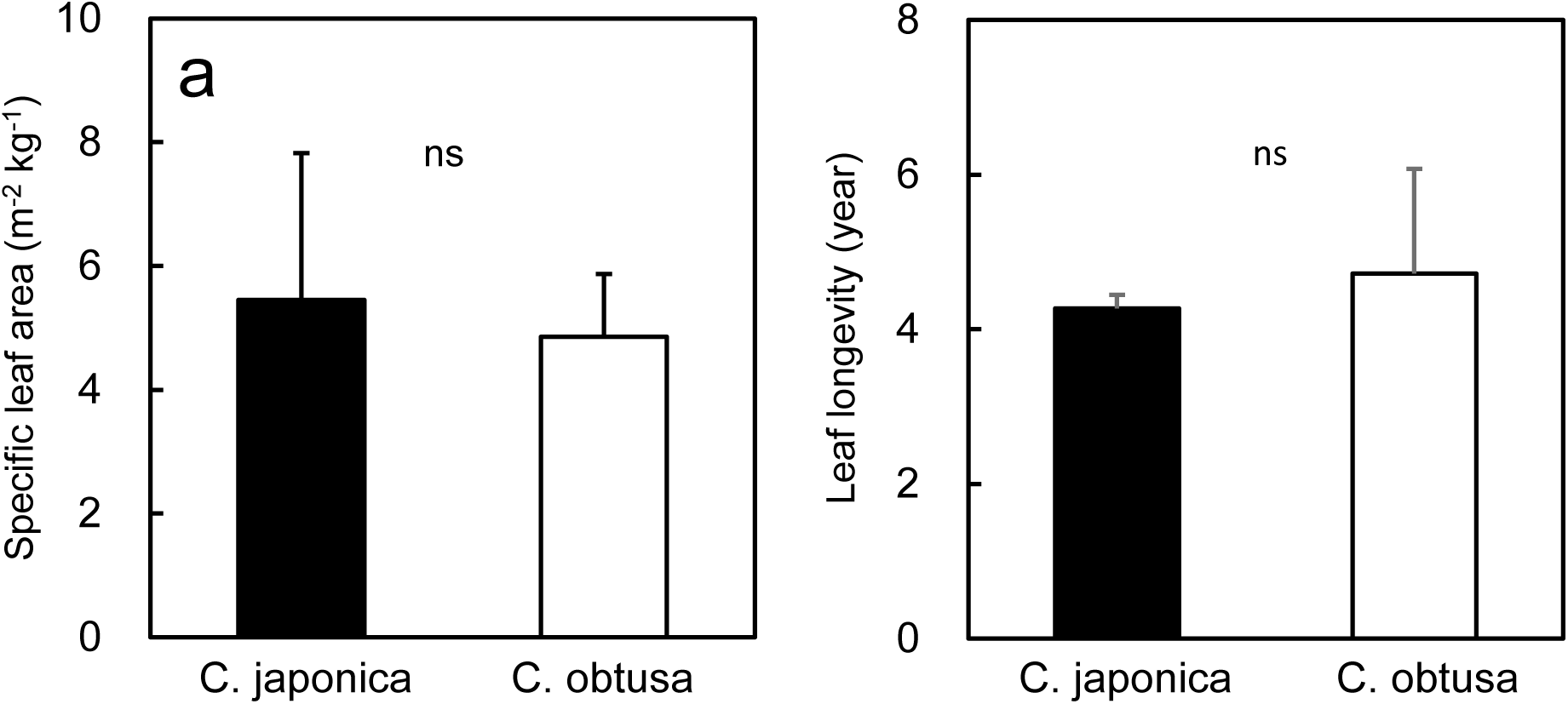
Specific leaf area and leaf longevity of *C*. *japonica* and *C*. *obtusa*. Differences between *C. japonica* and *C. obtusa* were subjected to a t-test; ***, P<0.001; xx, P<0.01; x, P<0.05; ns, not significant.

This is analogous to the effects that the different anatomy of sun/shade leaves has on foliar photosynthesis or the different structure of grass stands (steep foliage angle)/forb type (vertical foliage) on canopy photosynthesis (Monsi and Saeki, 1953; Terashima and Saeki, 1983; Terashima and Hikosaka, 1995). Similar to sun leaves with thick tissue layers and grass stands with high LAI, needle clumping would be favourable only if incident light is high and penetrates deep into the shoots. Under these conditions, whole shoot productivity could be higher than that of planar shoots where shoot photosynthesis saturates at lower light levels. If the incident light is low, however, light attenuates on the upper layers of the shoots without being transmitted to deeper layers. Under these circumstances, planar shoots are preferable for higher efficiency of weak light capture just as shade leaves or forb-type stands are preferred under low light availability. The differences in the light interception between the two shoot types were also reflected in photosynthetic light response curves. *C*. *japonica* had a lower initial slope (0.31) and convexity (0.59) than *C*. *obtusa* (initial slope, 0.48; convexity, 0.66) since photosynthesis increases and saturates at a slower rate with increasing irradiance in three-dimensional shoots (data from SugiHinokiDB). This is a well-documented pattern in the photosynthetic light response curves of sun/shade leaves and indicates that *C*. *obtusa* has characteristics of shade leaves in comparison with *C*. *japonica*. Although our hypothesis that photosynthetic capacity would be higher in *C*. *japonica* than in *C*. *obtusa* was not supported on a per needle basis, their adaptation to different light environments is more apparent in their light use and photosynthesis at the shoot level.

The two shoot types may also differ in total light interception per day. Light interception is the most efficient when leaves are oriented to face the direction of the light source. Therefore, planar shoots (*C*. *obtusa*) can intercept light more efficiently than shoots that have needles with various orientations (*C*. *japonica*) when the light source is just above them and is weak. However, since the solar azimuth angle changes considerably during a day, shoots that have needles with various orientations may be able to intercept more light and have higher assimilation rate on a daily basis than planar shoots, especially under strong light conditions (Stenberg et al., 1995; Stenberg, 1996). These results also imply that *C*. *japonica* is advantageous over *C*. *obtusa* in open habitats where strong light is available for longer times during a day. Such environments do appear in early stages of plantations before canopy closure. Thus, *C*. *japonica* may be able to grow faster than *C*. *obtusa* due to higher daily photosynthesis in this stage of a plantation.

#### 3.2.2 SLA

SLA (projected foliage area per foliage mass) was not significantly different between *C*. *japonica* and *C*. *obtusa* (Fig. 3a). This contradicts our hypothesis and the vast majority of studies that have reported a correlation between SLA and the relative growth rate across a wide range of plant species (Poorter and Remkes, 1990; Cornelissen et al., 1996; Atkin et al., 1998; Poorter and Van der Werf, 1998; Reich et al., 1998a; Wright and Westoby, 2000, Osone et al., 2008). From the viewpoint of growth analysis, SLA contributes to the relative growth rate because a high SLA is assumed to represent a large photosynthetic surface (=area of light interception) per given foliage biomass (Hunt, 1978; Lambers et al., 2008). However, as discussed above, the relationship between needle area and light interception is not straightforward for three-dimensional shoots, and it is possible that *C*. *japonica,* with shoots of various needle orientations, has a higher daily assimilation rate than *C*. *obtusa* with planar shoots in certain environments. On the other hand, the consistent SLA of the two species appears reasonable from the viewpoint of the leaf economic spectrum, a multivariable correlation between key chemical, structural and physiological properties of leaves based on the carbon and nitrogen economy (e.g., Wright et al., 2004). One of the key axes of the leaf economic spectrum is that SLA is positively correlated with the mass based photosynthetic capacity and foliage N concentration and is negatively correlated with foliage longevity (Kikuzawa and Lechowicz, 2011). If so, the two species with a similar photosynthetic capacity (on a needle basis), foliage N concentration (needle basis) and foliage longevity (Fig. 2), should also be similar in SLA.

#### 3.2.3 Biomass ratio between foliage and other organs

The biomass ratio between organs is also a factor that affects the growth rate because a relatively high foliar mass equates to a relatively large photosynthetic organ if other factors are equal (Hunt 1978; Lambers et al. 2008). At a given DBH or total biomass, the stem and root biomasses were not significantly different between the species, but foliar biomass and therefore the foliar mass ratio (LMR) were higher in *C*. *japonica* than in *C*. *obtusa* (Fig. 4). This tendency was more pronounced at smaller DBHs or total biomass, suggesting that the initial greater LMR could contribute to faster growth at early stages of *C*. *japonica* growth. There could be several reasons for the higher LMR in *C*. *japonica*. Since plants grown under high soil resource availability allocate more biomass to foliage at the expense of roots (Osone and Tateno, 2005, and references therein), *C*. *japonica*, which is often planted in fertile wet sites, might allocate more biomass to foliage than *C*. *obtusa*, which is planted in less fertile and dry sites. Another possible reason is related to the shoot and crown form of *C*. *japonica*. Generally, the leaf area index (≈leaf biomass) is larger in canopies where the light absorption coefficient (*k*) is smaller because light penetrates deeper into the canopy (I_x_ = I_0_ exp(-*k*F_x_, where I_x_ and I_0_ are the photon flux density at depth x and outside the canopy, respectively) and Fx is the cumulative LAI at depth x, first proposed by Monsi and Saeki 1953). One of the factors that decreases *k* is a steep foliage inclination (Terashima and Hikosaka, 1995 and references therein), and *C*. *japonica,* which has three-dimensional shoots, had a lower *k* (0.38) than *C*. *obtusa* with planar shoots (0.97, but only one data entry was available). Therefore, the shoot morphology that allows light penetration deeper into the crown may cause *C*. *japonica* to have a thick canopy (=high LMR).

**Fig. 4.**
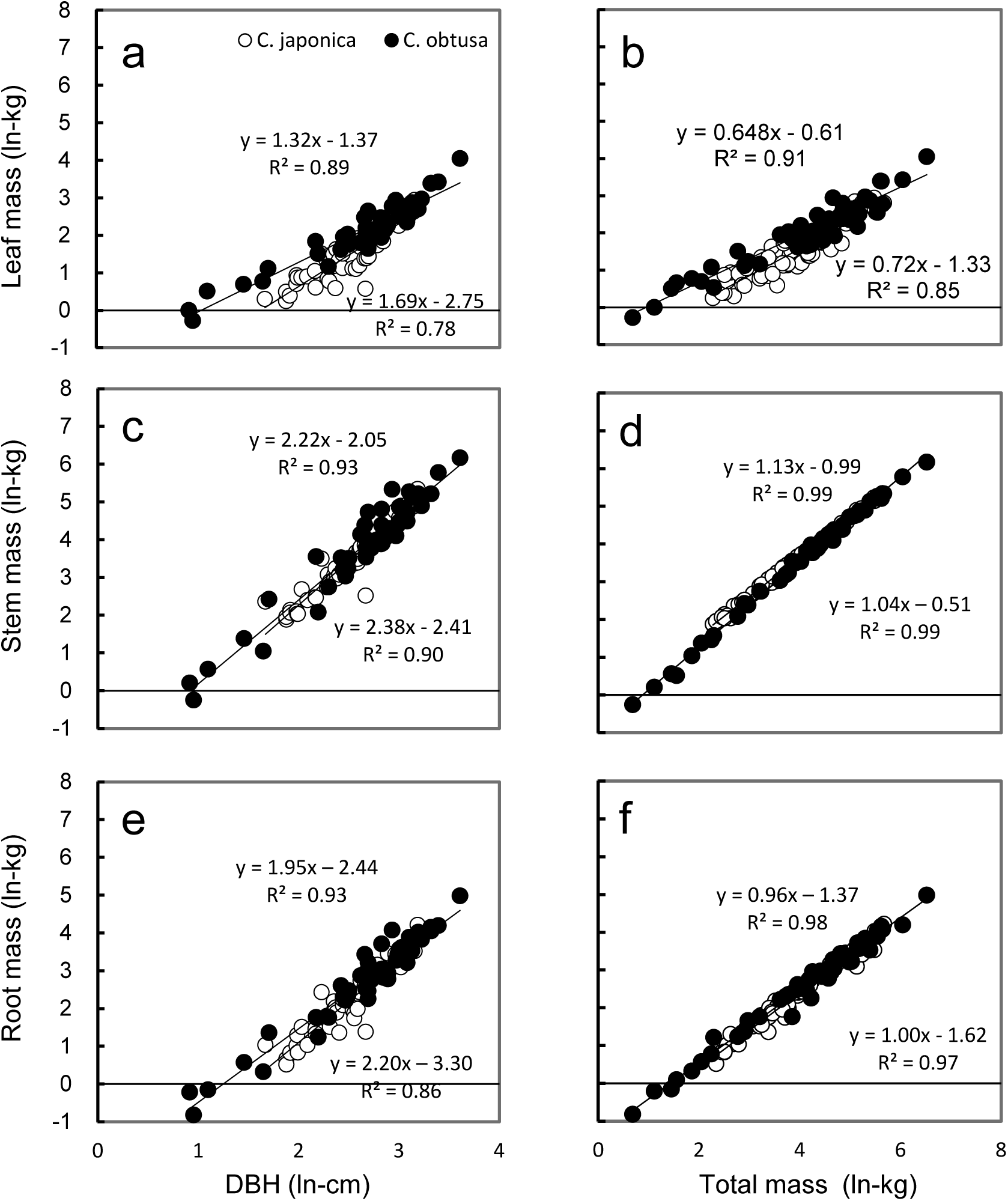
Ln-transformed organ biomass in relation to ln-transformed DBH (a, c, e) and ln-transformed total mass (b, d, f).

### 3.3 Differences in traits related to water use

#### 3.3.1 Stomatal conductance and transpiration rate

*C*. *japonica* exhibited 1.7- and 1.5-fold higher stomatal conductance to CO_2_ (gs_an_) and transpiration rate (E_an_), respectively, compared with *C. obtusa* (Fig. 5a, 5b). In concert with this, stomatal distribution and anatomy differed largely between the species. Coniferous stomata are generally distributed unevenly on foliar surfaces, forming a species-specific pattern of stomatal clusters called a “white band”. *C. japonica* has two thick white bands on every surface of the triangular pyramid-shaped needles, whereas *C. obtusa* has y-shaped white bands along the rims on the abaxial surface of its scales (Ma et al., 2007; Yazaki et al., 2015; Kim, 2018). In addition to this larger proportional area of white bands to the total foliage surface, the stomatal diameter of *C*. *japonica* was 1.5 times higher than that of *C*. *obtusa* (Fig. 5c).

**Fig. 5.**
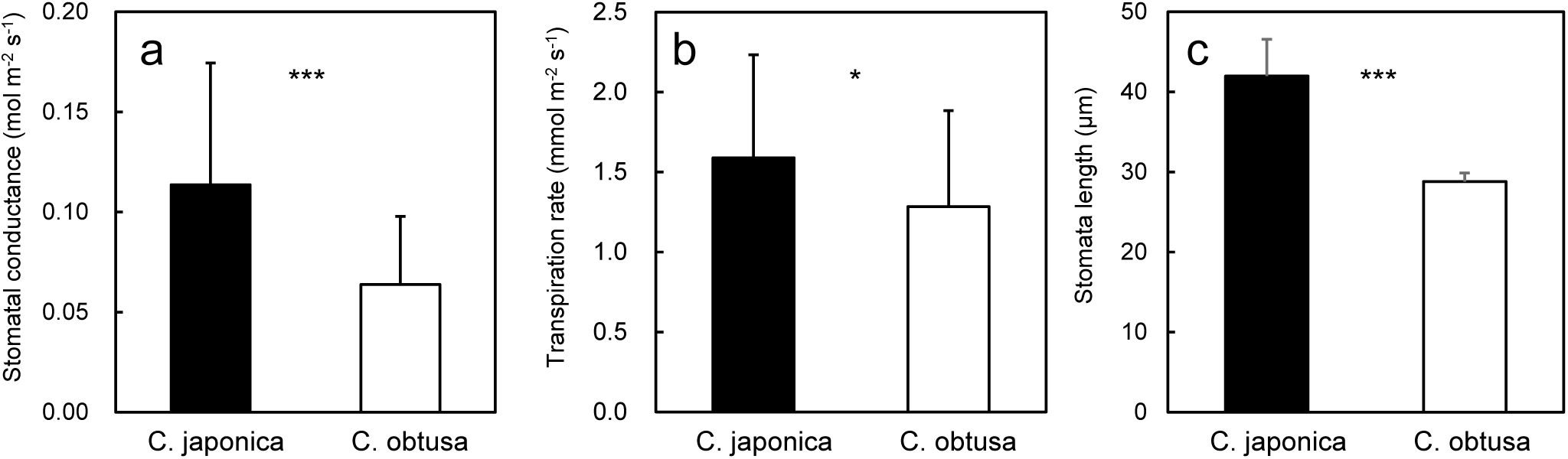
Leaf traits of *C. japonica* and *C*. *obtusa*. In *C*. *japonica*, each trait is presented on a shoot silhouette area basis and projected needle area basis. Differences between each *C. japonica* trait (shoot or needle basis) and each *C*. *obtusa* trait were determined by a t-test; ***, P<0.001; xx, P<0.01; x, P<0.05; ns, not significant.

Despite the lower gs_an_ and E_an_ in *C*. *obtusa*, Amax_an_ was not significantly different between the species (Fig. 2a), suggesting that water use efficiency, that is, gs_an_ or E_an_ divided by Amax_an_, is higher in *C*. *obtusa* than in *C*. *japonica*. The lower gs_an_ and E_an_ and higher water use efficiency are regarded as adaptations of drought-tolerant plants (Kozlowski and Pallardy, 2002), which supports the hypothesis that *C. obtusa* is more drought tolerant than *C. japonica*. However, adaptation to environments with different levels of water availability is represented not only by potential gas exchange rates but also by the magnitude and speed of the stomatal response to changing environments (Oren et al., 1999; Meinzer et al., 2016). Although stomatal sensitivity are less often measured than potential gas exchange rates are and they are not collected in SugiHinokiDB, some studies that estimated canopy stomatal conductance by sap flow measurements suggested that mature *C. japonica* trees were less sensitive to increasing VPD than *C. obtusa* (Kumagai et al., 2008; Tsuruta et al., 2020).

#### 3.3.2 Foliage PV curve parameters

Among the parameters of the pressure-volume curve, Ψ_tlp_ (foliage water potential at the turgor loss point) showed the clearest species difference: Ψ_tlp_ was larger in *C*. *obtusa* than in *C*. *japonica* throughout the year (Fig. 6a). Since plants with more negative Ψ_tlp_ are able to maintain cell turgor pressure under drought stress, thereby sustaining stomatal conductance, photosynthesis and growth, Ψ_tlp_ is thought to be predictive of the drought tolerance of a plant species (Scholander et al., 1965; Tyree and Jarvis, 1982; Tognetti et al., 2000; Baltzer et al., 2008; Blackman et al., 2010; Zhu et al., 2018). Indeed, midday foliage water potential was lower in *C*. *obtusa* than in *C*. *japonica* during summer (Fig. 6c) indicating that *C*. *obtusa* continued to open stomata until the water potential more decreased than that of *C*. *japonica.* This supported the hypothesis that *C*. *obtusa* is more drought tolerant than *C*. *japonica*. There are three possible ways in which Ψtlp becomes more negative: the accumulation of solutes such as sugar (decreases Ψπsat), a reduction in the symplastic water content through the redistribution of more water outside the cell walls (decreases Ψπsat) and an increase in cell wall flexibility (decreases ℇ) (Bartlett et al., 2012). Ψπsat was slightly lower in *C*. *obtusa* than in *C*. *japonica*, but the species difference was marginal throughout the year (Fig. 6b). Unfortunately, ℇ values were limited in the database, and we could not make reliable comparisons of monthly ℇ between the species, but the small difference in Ψπsat suggests that the lower Ψtlp of *C*. *obtusa* might be due to its presumed smaller ℇ. Species differences in Ψtlp are more often correlated with Ψπsat than with ℇ since a unit decrease in Ψπsat causes a larger decrease in Ψtlp than in ℇ (Saito et al., 2003; Mitchell et al., 2008; Bartlett et al., 2012; Meinzer et al., 2016). However, as was shown in the present study, there are also studies that showed a correlation between Ψtlp and ℇ (Niinemets, 2001).

**Fig. 6.**
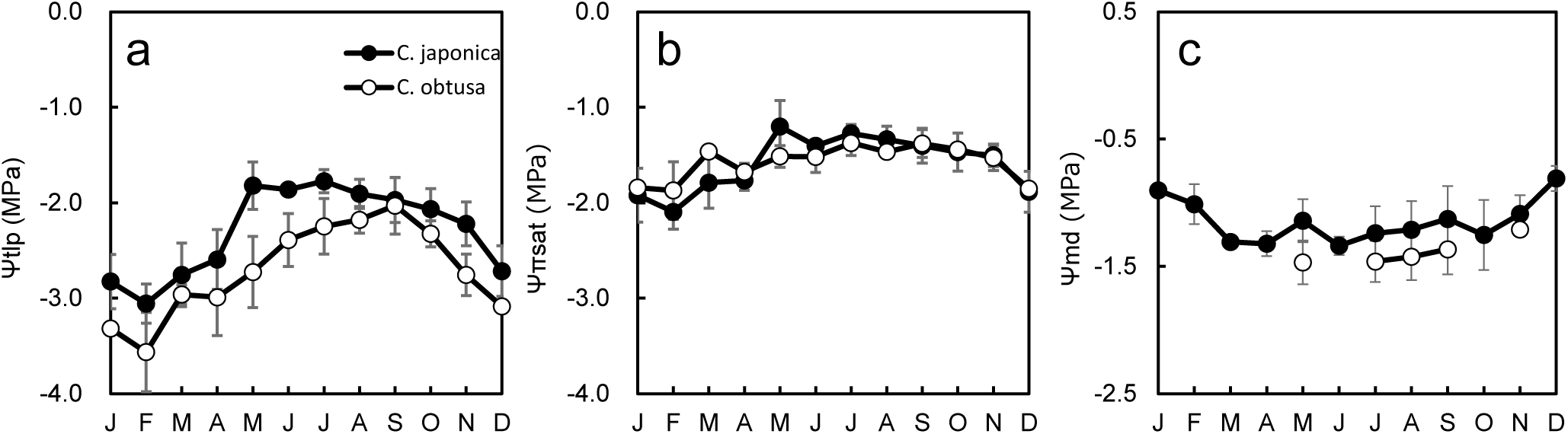
Seasonal changes in pressure volume parameters and midday leaf water potential.

The other marked difference in the PV curve parameters was that *C*. *japonica* had higher ℇ in winter (Nov-Apr) compared to summer (May-Oct), whereas *C*. *obtusa* showed no seasonal changes in ℇ (Fig. 7a). Winter increases in ℇ were also reported for *Eucalyptus* species (Valentini et al., 2012), Taiwan cedar (Hara et al., 1998) and Patagonian woody shrubs (Scholz et al., 2011; Zhang et al., 2016) and are thought to reduce physical injury to cell membranes by making cell walls more rigid (higher ℇ) during extracellular freezing and/or thawing processes. However, if increases in ℇ occur in response to freezing resistance, it is not clear why only *C*. *japonica* exhibited this response while both species exhibited a winter decrease in Ψπsat (Fig. 7), which is also a well-known response to freezing (Sakai, 1982; Sakai and Larcher, 1987). Considering that *C*. *japonica* is more water demanding, another explanation may be possible for the winter increase in ℇ — the ‘cell water conservation hypothesis’ (Cheung et al., 1975; Bartlett et al., 2012). Theoretically, reductions in Ψπsat decrease both Ψtlp and RWCtlp. However, a coordinated reduction in Ψπsat and an increase in ℇ would lower Ψtlp while maintaining a constant RWCtlp, which would result in tolerance for freezing and also prevent dangerous cell dehydration and shrinkage. In *C. japonica*, ℇ was negatively and strongly correlated with Ψπsat (Fig. 7a), and in accordance with the theory, RWCtlp remained constant irrespective of Ψπsat (Fig. 7b). In contrast, in *C. obtusa*, where reductions in Ψπsat did not accompany increases in ℇ, RWCtlp decreased with reductions in Ψπsat (Fig. 7a, 7b). Plant species have a minimum cellular water content to maintain metabolic functions (Lawlor and Cornic, 2002). Since *C*. *japonica* has a higher minimum tissue water requirement for survival than *C*. *obtusa* (Satoo, 1956), it may adjust ℇ in coordination with Ψπsat to constantly maintain RWCtlp above this minimum tissue water content.

**Fig. 7.**
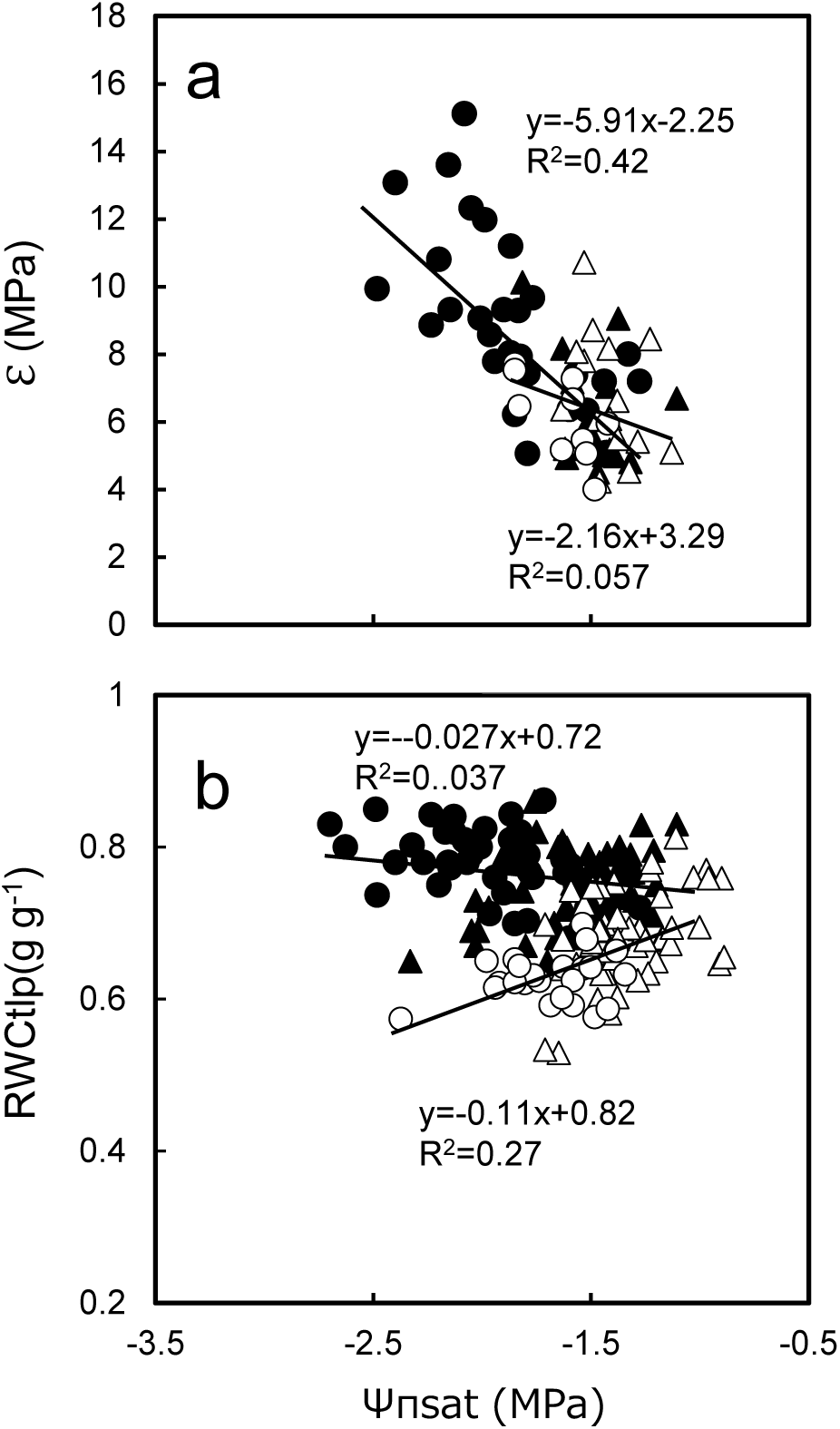
Relationship between pressure-volume parameters. Data measured from May-Oct and Nov-Apr were pooled. Filled triangle, *C. japonica* data from May-Oct; Filled circle, *C. japonica* data from Nov-Apr; Open triangle, *C. obtusa* data from May-Oct; Open circle, *C. obtusa* data from Nov-Apr.

#### 3.3.3 Hydraulic architecture

Soil-to-foliage hydraulic conductance (K_S-L_), an index of whole-plant hydraulic efficiency, was 1.6 times higher in seedlings of *C*. *japonica* than in seedlings of *C*. *obtusa* (Fig. 8a). Hydraulic conductance is a major determinant of plant water status and stomatal behaviour (Zimmermann, 1978; Meinzer et al., 2001) because of the relationship E_an_ = K_S-L_ (Ψ_pd_-Ψ_L_), where Ψ_pd_ is the foliage water potential measured at predawn, which represents the soil water potential, and Ψ_L_ is the foliage water potential. In accordance with this relationship, *C. japonica*, with a higher K_S-L_, had a higher transpiration rate than *C. obtusa* (Fig. 5b), maintaining a higher midday foliage water potential (Fig. 6c). Extensive measurements of K_S-L_ across species have revealed the adaptive significance of hydraulic conductance across functional groups; that is, pioneer, mesic and drought-avoiding species have higher hydraulic conductance than late successional, xeric and drought-tolerant species (Tyree et al., 1998, Nardini and Tyree, 1999, Nardini and Pitt, 1999; Stratton et al., 2000, references in Meinzer et al., 2001). The higher K_S-L_ in *C*. *japonica* than in *C. obtusa* appears in line with this. However, since K_S-L_ depends on the water transport distance, here, we used only seedling data where comparisons at similar sizes (ca. 1 m in height) were possible and are not sure whether this is also true for adult trees.

**Fig. 8.**
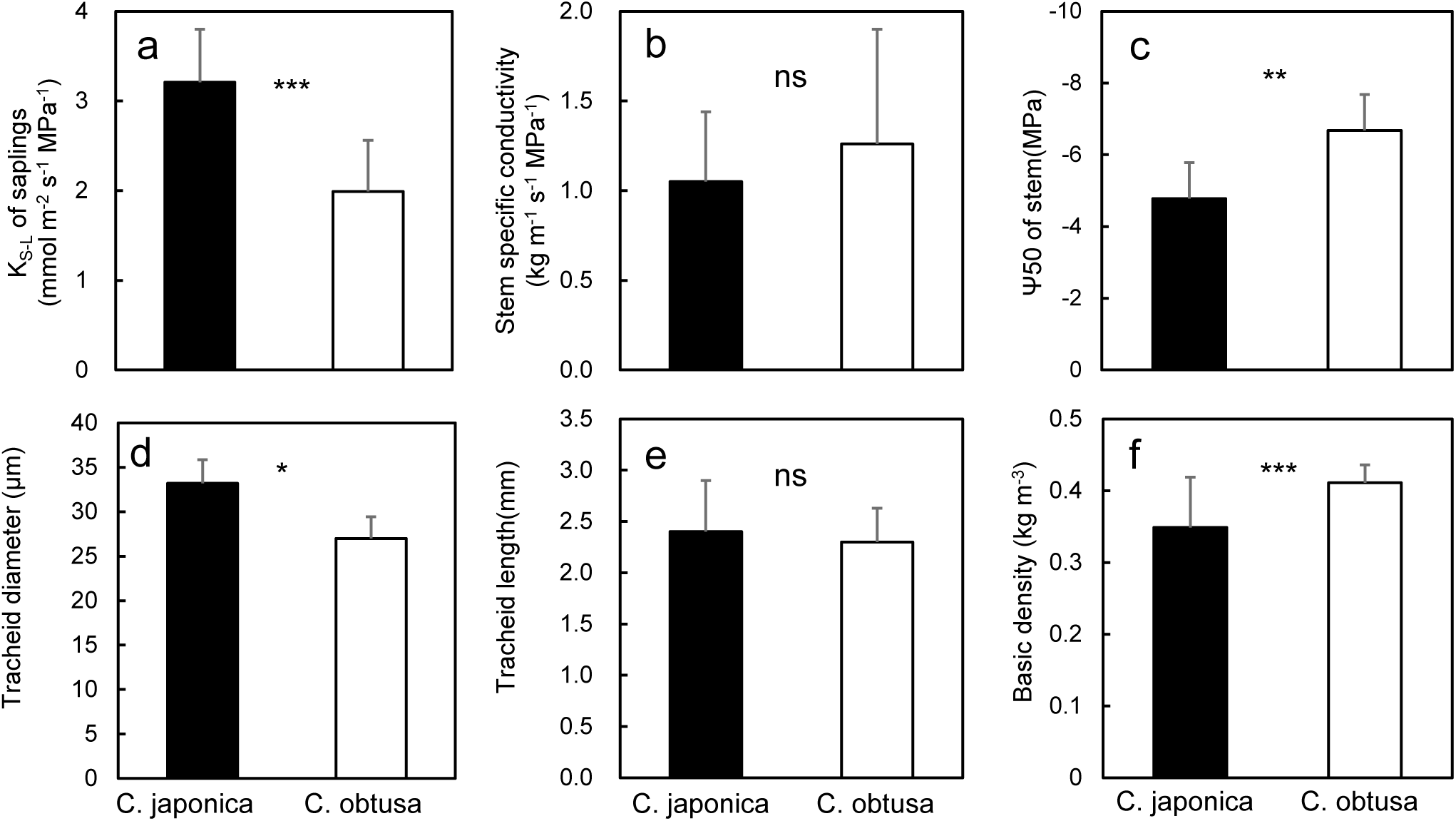
Stem hydraulics and xylem anatomy. Differences between the species were examined by a t-test; ***, P<0.001; xx, P<0.01; x, P<0.05; ns, not significant.

Since K_S-L_ is determined by the conductivity of all organs along the whole water transport pathway, knowledge on the hydraulic conductivity of each would provide more insight for water use. Unfortunately, we still have limited knowledge on organ hydraulic conductivity of these species. The only available information, i.e., stem-specific conductivity presented as hydraulic conductance per stem sapwood area (K_stem_), was not significantly different between the two species (Fig. 8b), suggesting that the higher K_S-L_ of *C*. *japonica* could be due to the higher foliage and/or root hydraulic conductivity of this species.

In addition to hydraulic efficiency, water transport safety is another important factor of hydraulic architecture. Although we have even less information for hydraulic safety than for hydraulic conductivity, xylem pressure corresponding to a 50% loss of conductivity (Ψ50), an index of safety from embolism (Choat et al., 2012), was 1.4 times higher in *C*. *obtusa* than in *C. japonica* (Fig. 8C). This means that *C*. *obtusa* can endure more severe negative pressure than *C*. *japonica*, which supports the empirical knowledge that this species is more tolerant to drought. However, in Japan, where precipitation is generally high throughout the year, plants rarely experience such extreme negative pressure, represented by the Ψ50 (−6.7 MPa) in the xylem of *C. obtusa*, and the extent to which the differences in Ψ50 are relevant to their habitat preference is not obvious. Recently, Ψe, which is the xylem pressure at the start of conductivity loss, has been considered to be a more suitable index for drought tolerance than Ψ50 in nonextreme habitats (Meinzer et al., 2009). Although Ψe is less focused on than Ψ50, studies show that plants control stomatal conductance to maintain xylem pressure near Ψe (Sparks and Black, 1999; Brodribb et al., 2003; Domec et al., 2008), indicating that Ψe could be a key factor linking stomatal control and xylem pressure. Measurements of Ψe, therefore, would lead to a better understanding of differences in the drought response of these species. In conifers, hydraulic vulnerability segmentation, that is, a lower resistance to embolism in the distal segments that ensures the safety of the more proximal stems, is also a common hydraulic strategy (Johnson et al., 2016). In this case, not the hydraulic resistance of stems themselves but the differences in resistance between distal segments and more proximal segments (e.g., differences in Ψ50 between leaves and stems) should be more important. However, because characterization of hydraulic safety is tedious and time-consuming, relatively few data are available in this field overall (Blackman et al., 2010; Santiago et al., 2018), and this is also the case for these species (Table 1).

The tracheid diameter of the stem xylem was 1.26 times larger in *C*. *japonica* than in *C*. *obtusa*, and the length was not significantly different (Fig. 8d, 8e). The basic density of stem wood (wood density) was lower in *C*. *japonica* than in *C*. *obtusa* (Fig. 8f). Generally, the tracheid structure is considered to be closely related to water transport efficiency and drought safety, such as cavitation in the xylem (Delzon et al., 2010; Jansen et al., 2012). Conducting efficiency increases with tracheid diameter according to the Hagen–Poiseuille law, and it is also correlated with the tracheid length because a longer tracheid can result in conductive pits in the end walls, where water flow is significantly limited, being farther apart (Hacke et al., 2006; Pittermann et al., 2006a, 2010). If so, stem specific conductivity should be higher in *C*. *japonica* than in *C*. *obtusa*, but it was not significantly different between the species (Fig. 8b). The reason is not clear, but the pit structure, which could affect conductivity and for which we have no information for these species, might play a role (Delzon et al., 2010; Jansen et al., 2012). A more negative Ψ50 is also known to be associated with a smaller tracheid diameter and greater basic density because it requires mechanical strength to support the xylem conduit against implosion caused by negative pressures (Hacke et al., 2001; Pittermann et al., 2006b; Ogasa et al., 2013). The smaller tracheid diameter, higher wood density and more negative xylem Ψ50 in *C*. *obtusa* compared with *C*. *japonica* are in line with the hypothesis.

### 3.4 Age and height dependency

Tree age and/or size, especially height, usually affect many foliar functional traits, such as photosynthetic traits, stomatal behaviour, morphology, water use and nutrient concentrations, in various tree species, including angiosperms and gymnosperms in tropical, temperate, boreal and even semiarid areas (Ellsworth and Reich, 1993; Ryan and Yoder, 1997; Bond, 2000; Rijkers et al., 2000; Koch et al., 2004; Duursma et al., 2006; Kenzo et al., 2006, 2015, 2016; Ryan et al., 2006; Azuma et al., 2016; Chin and Sillett, 2019; Liu and Hikosaka, 2020). These age- and/or size-related foliage changes are important for understanding the forest growth rate, timber yield and carbon balance because they have a strong relationship with individual-based functional traits such as photosynthetic productivity and drought tolerance (Ryan et al., 2006). In this section, we demonstrate the effects of tree age and/or height on foliar nutrient concentrations, especially nitrogen (N), phosphorus (P) and potassium (K), specific leaf area (SLA) and midday minimum water potential in the foliage, using a database. These foliar traits generally indicate the photosynthetic ability and stomatal behaviour of plants, e.g., there is a positive relationship between the nitrogen concentration and maximum photosynthetic rate at light saturation, and the minimum midday water potential indicates plant drought stress (Kramer and Boyer, 1995; Lamberts et al., 1998). If we can clearly demonstrate those relationships in *C*. *japonica* and *C*. *obtusa*, the findings will contribute to the development of accurate estimation models to predict future forest production and carbon dynamics in those forests. In addition, since the analysed data sets were obtained from sunlit foliage, the effects of light intensity on tree height can be ignored as much as possible.

#### 3.4.1 Foliage nutrient contents

Changes in foliar nutrient concentrations with tree age and/or height showed different patterns depending on tree species and nutrient type. The foliar N concentration was similar among the age classes of *C*. *japonica*, whereas that of *C*. *obtusa* decreased significantly with age (Fig 9a, 9b). On the other hand, foliar P and K concentrations in *C*. *japonica* decreased significantly with age class (Fig. 9c, 9e). A similar age-dependent reduction in the P concentration was observed in *C*. *obtusa* (Fig. 9d), although the K concentration was constant with age (Fig. 9f). Various patterns of changes in foliar nutrient contents with tree size and/or age have been reported. Although several cases indicate an increase in the foliar nitrogen content with tree size and age (Chen et al. 2018 for N of Fabaceae tree species), many other cases showed decreased (Beets and Madwick, 1988; Polglase and Attiwill, 1992; Bargali et al., 1992; Hooker and Compton, 2003; Tanaka-Oda et al., 2010; Yang and Luo, 2011) or constant (Polglase and Attiwill, 1992; Clinton et al., 2002; Kenzo et al., 2012, 2015; Givnish et al., 2014; Chen et al., 2018) contents of foliar N and P with tree size and age. These variations may be caused by species-specific traits, soil nutrient availability, and sampling effects caused by rather small datasets (Thomas and Winner, 2002). In the present study, it is believed that the influence of sampling effects is small due to the large data sets, e.g., 200-1500 data points (forest age up to 80 years old, tree height up to 40 m and grown on various forest soil types), used to examine the relationship between each tree species and nutrient. Increasing nutrient accumulation to living biomass and coarse woody debris with forest development may cause decreased foliar nutrient concentrations with tree age and height through a reduction in soil nutrient availability (Bond, 2000; Clinton et al., 2002; Chen et al., 2018). A reduction in soil nutrient concentrations with forest development has been reported in the early stages of forest growth (Ohta, 1990; Kleinman et al., 1995; Hattori et al., 2013). In addition, higher drought stress, such as hydraulic limitation with tree height, promotes the investment of carbon into foliage compared with physiological functions, which require more N, P, and K to protect against dehydration (Niinemets, 1997; Thomas, 2010; Kenzo et al., 2012). As a result, invested carbon dilutes the foliar nutrient concentrations. These changes are likely to be true for changes in the P and K concentrations of *C*. *japonica* and *C*. *obtusa* trees. In contrast, several authors have suggested that tree size- or age-related foliar N contents may have a unimodal relationship rather than a simple linear relationship (Ishii et al., 2008; Thomas, 2010; Kenzo et al., 2012). If nonlinear changes in foliar N occurred in each *C*. *japonica* and *C*. *obtusa* stand, it may be difficult to identify a clear linear relationship of N concentration with tree size and height by using the present pooled analysis. Further studies should consider other factors, such as soil type and slope position, to understand more detailed changes in foliar nutrients with age and height.

**Fig. 9.**
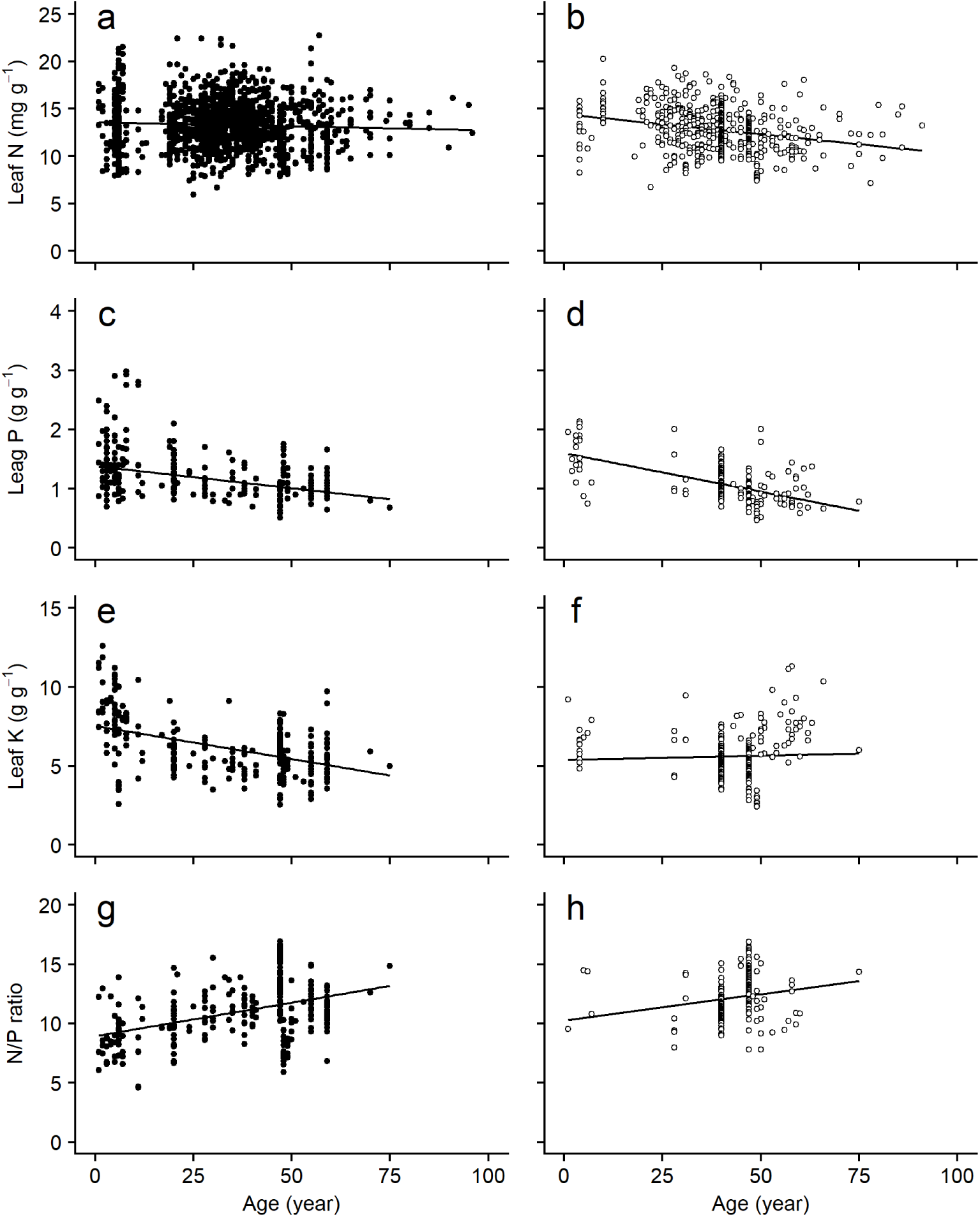
Changes in leaf nutrient contents with tree age. Data from plants grown under experimental conditions or in pots were excluded. Data from plants younger than 100 years old are shown.

**Fig. 10.**
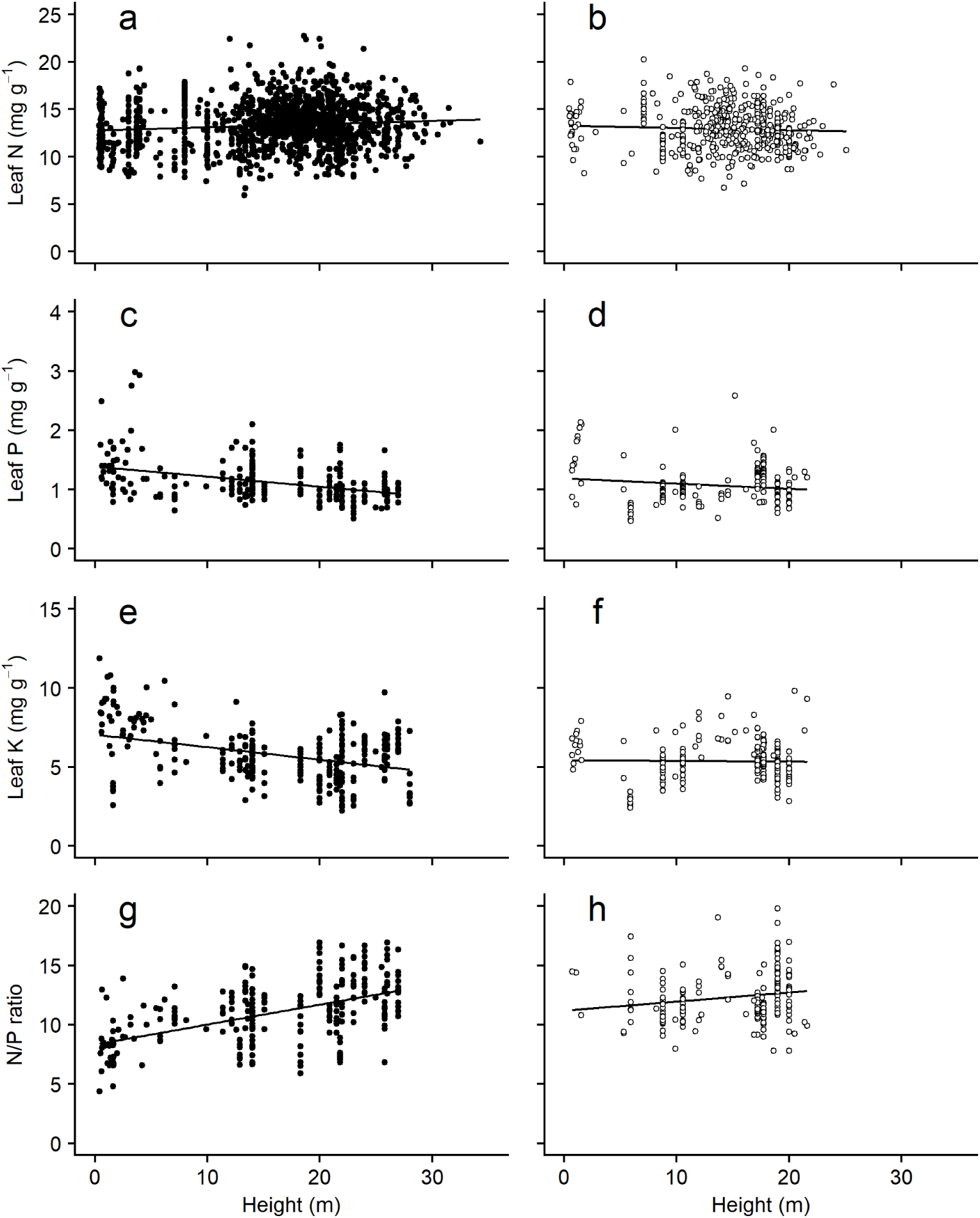
Changes in leaf nutrient contents with tree height. Data from plants grown under experimental conditions or in pots were excluded.

The difference in the changes in each nutrient between *C*. *japonica* and *C*. *obtusa* is believed to reflect the interspecific differences in nutrient demand and usage and growth habitats. For example, it has long been known that *C*. *japonica* forests are planted under moister conditions in nutrient-rich soil, and their nutrient dynamics are faster than those of *C*. *obtusa* forests, which usually grow on upper slopes with poor nutrient and water availability (e.g., Sawata and Kato, 1993; Tanikawa et al., 2014). These differences may affect the age- and height-dependent changes in foliar nutrients.

The stoichiometry of N to P changed with tree age and height based on different patterns of changes in foliar N and P concentrations. In *C*. *japonica*, where the foliar P concentration decreased with forest age and the N concentration did not change, the NP ratio increased significantly with the age of the forest (Fig. 9g). In *C*. *obtusa*, the N:P ratio tended to increase but was not significant (slope = 0.024, P = 0.34). The NP ratio is an index of soil nutrient limitation, e.g., P limitation occurs if the ratio is higher than 16, N limitation typically occurs if the ratio is lower than 14, and N and P are co-limiting if the value is 14-16 (Koerselman and Meuleman, 1996). The maximum NP ratio of *C*. *japonica* is 16.9, and 87.2% of all data points have values of 14 or lower, suggesting that *C*. *japonica* stands are generally N-limited. However, since the NP ratio increases with age, the stands shift from being N-to being P-limited with maturity. *C*. *obtusa* also has an NP ratio of 14 or less, and approximately 77% of the data points are considered to be in the N-limited range (Fig. 9h).

#### 3.4.2 Foliage water potential and specific leaf area (SLA)

As tree height increases, drought stress increases in the upper part of the canopy, which in turn affects morphological and physiological responses of tree foliage (Koch et al., 2004; Ryan et al., 2006). The foliar midday water potential, which indicates the degree of tree drought stress, significantly decreased with height and age in *C*. *japonica* trees (Fig. 11a, 12a) and with age in *C*. *obtusa* trees (Fig. 11b). These reductions in tree height have been reported for various tree species worldwide and cause hydraulic limitations, such as a reduction in photosynthesis through stomatal limitations (Fredericksen et al., 1996; Ryan and Yoder, 1997; McDowell et al., 2002; Woodruff et al., 2004; Ishii et al., 2008; Ambrose et al., 2009). Interestingly, the recovery of photosynthetic ability by grafting canopy tree shoots onto saplings of *C*. *japonica* supported the occurrence of hydraulic limitations in tall trees of this species (Matsuzaki et al., 2005). The slopes of Ψmd with the tree height of *C*. *japonica* and *C*. *obtusa* are −0.137 and −0.089 MPa m^-1^, respectively. These slopes are gentler than those of tropical rainforest trees (−0.0282 MPa m^-1^, Kenzo et al. 2015) and similar to the temperate conifer *Sequoia sempervirens* (−0.0100 to −0.0113 MPa m^-1^) and Douglas fir (−0.019 MPa m^-1^) (Koch et al., 2004; Ishii et al., 2008). In general, to tolerate a decrease in Ψ_md_, the foliar water potential at the turgor loss point (Ψ_tlp_) must be reduced (Inoue et al., 2017; Zhu et al., 2018). Foliar osmotic adjustment as well as structural strength to withstand low negative pressure to achieve lower Ψ_tlp_ and foliage strength is accompanied by a decrease in SLA. In fact, the SLA of sugi, whose Ψ_md_ significantly decreased with tree age and height, decreased with tree height and age. Many studies have reported that SLA decreases with tree age and height (Thomas and Winner, 2002; Mediavilla and Escuderl, 2003; Koch et al., 2004; Kenzo et al., 2006, 2015, 2016; Coble et al., 2014; Azuma et al., 2016). In addition, Thomas and Winner (2002) showed that SLA decreases with tree age in all existing studies based on meta-analysis. On the other hand, the SLA of *C*. *obtusa* did not show a significant change with either tree age or height (Fig. 11d, 12d). Although there was no clear reason, *C*. *obtusa* is more resistant to drought stress than *C*. *japonica,* and thus, changes in SLA may be small. However, the smaller size and age range in *C*. *obtusa* compared with *C*. *japonica* may cause this constant change in SLA because Shiraki et al. (2016) recently reported that SLA decreased significantly at a single canopy height in a *C*. *obtusa* tree. Thus, further data collection on older and taller *C*. *obtusa* trees is needed to understand tree age and size dependency with respect to SLA.

**Fig. 11.**
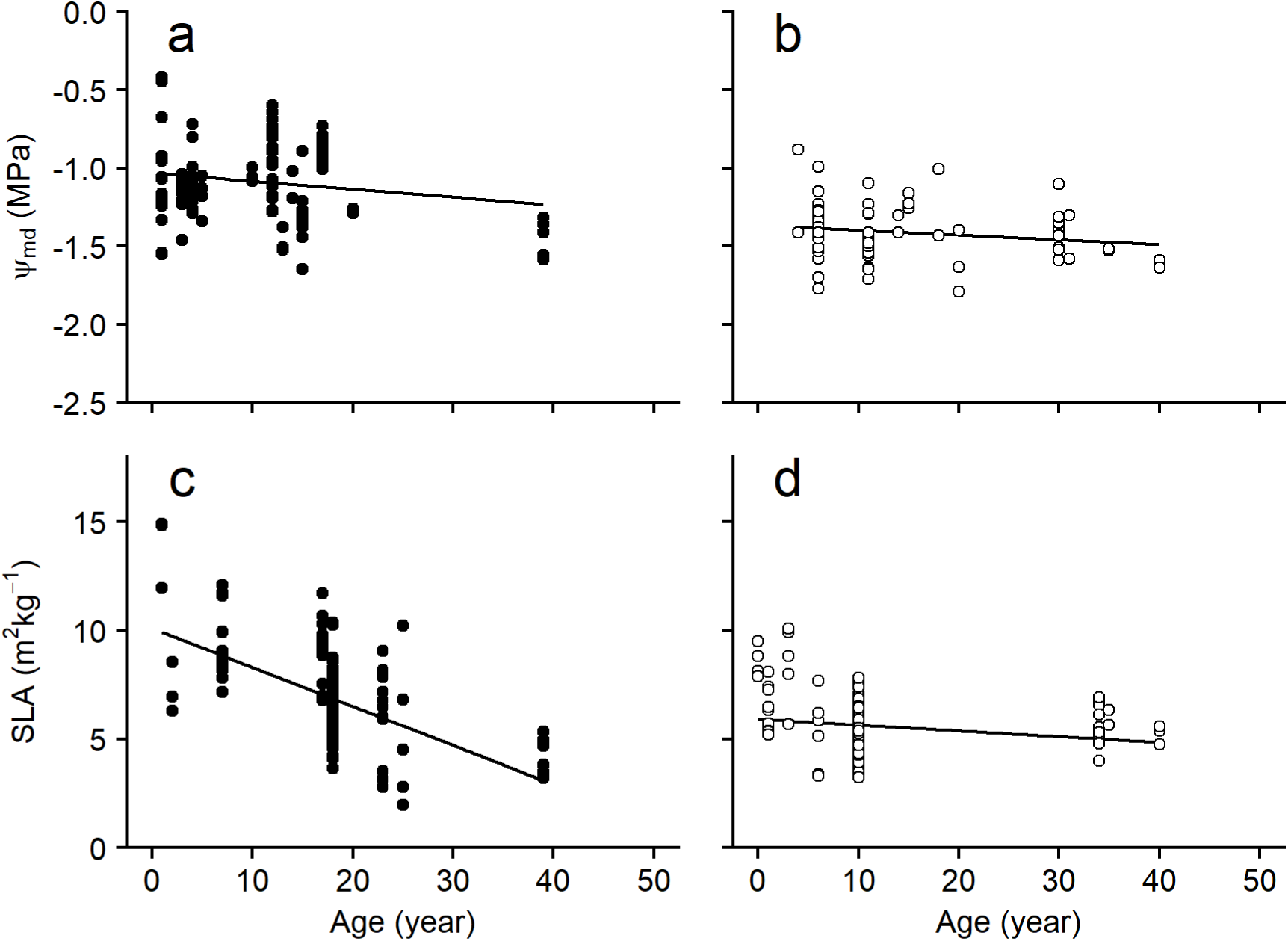
Changes in midday leaf water potential (Ψmd) and SLA with tree age. Data from plants grown under experimental conditions or in pots were excluded. Data from plants younger than 100 years old are shown.

**Fig. 12.**
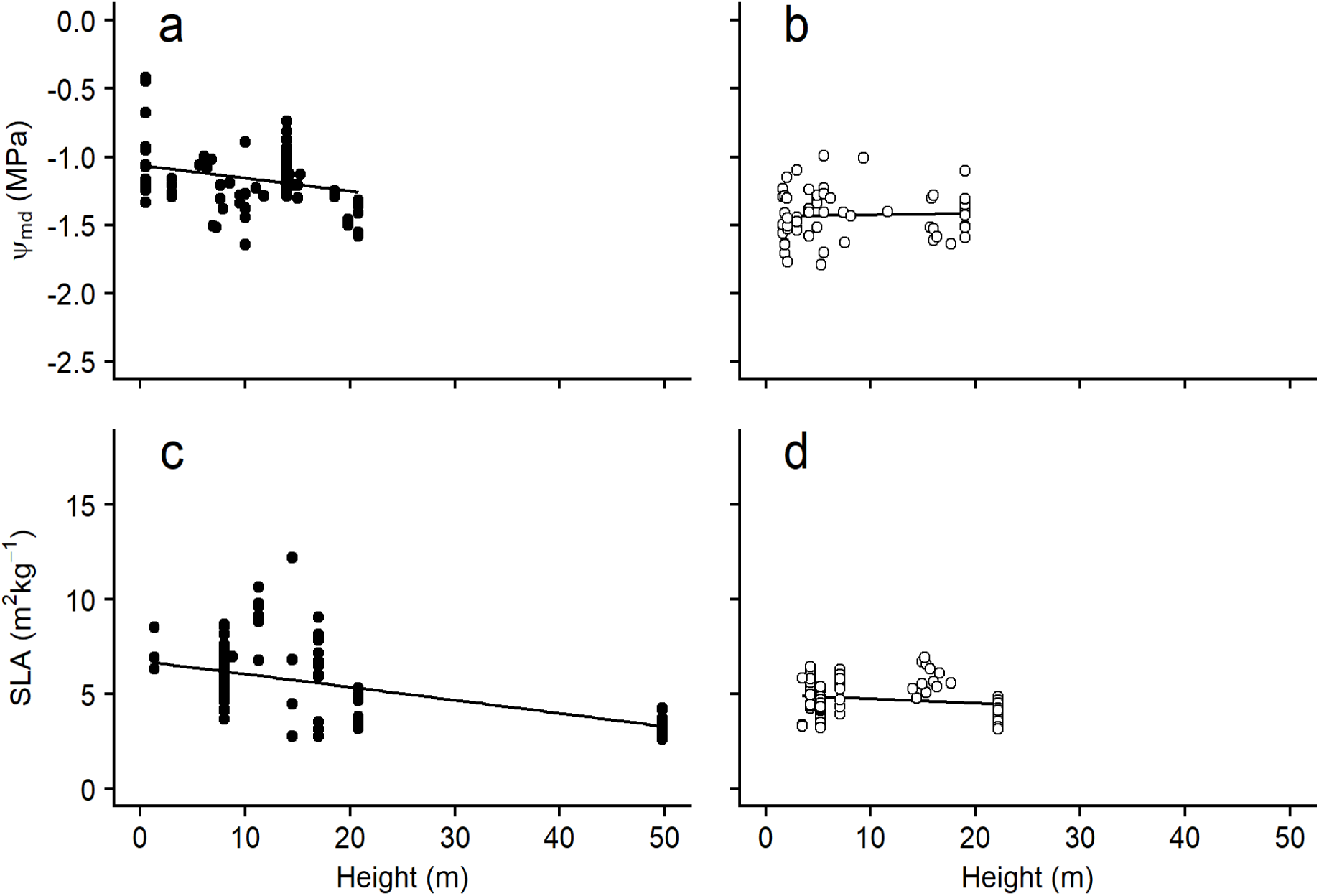
Changes in midday leaf water potential (Ψmd) and SLA with tree height. Data from plants grown under experimental conditions or in pots were excluded.

### 3.5 Comparison with broader species in *Cupressaceae* with respect to drought tolerance

Through the present analysis, clear differences in growth and water use characteristics were observed between *C*. *japonica* and *C*. *obtusa*, although the differences were at most ca. 50% (excluding stomatal size). The Cupressaceae family, to which *C*. *japonica* and *C*. *obtusa* belong, consists of more than 100 species with a marked diversity in physiology, morphology and habitat preference (Pittermen et al., 2012). How much do the contrasts we found between *C. japonica* and *C. obtusa* in the present study account for the ranges in traits exhibited by all Cupressaceae species? Here, we compare traits related to drought tolerance between the two species and other Cupressaceae species using data from Pitterman et al. (2012) to gain more insight into the ecological characteristics of these species.

Cupressaceae species first appeared in the warm and humid Mesozoic and differentiated in the cool and dry Cenozoic (Giffored and Foster, 1989). Reflecting the climatic conditions under which each species evolves, early diverging species prefer mesic-hydric habitats, while derived species are adapted to arid climates (Pittermann et al., 2012). As a result of these adaptations, the species exhibit a gradient of foliage morphology (from needle-like in basal species to scaly, similar to foliage in derived species) and other drought tolerance or water use characteristics (Pittermann et al., 2010; Brodribb et al., 2014; Gleason et al., 2016). In fact, *Juniperus sabina*, which has needle and scaly foliage at the ontogenic stage within the same individual, showed that needle foliage had higher photosynthesis with lower drought tolerance than scale-like foliage (Tanaka-Oda et al., 2010).

There was a significant relationship between Ψ_50_ and the foliar and xylem functional traits (Fig. 13). Species with higher wood density (Fig. 13a), lower xylem specific conductivity (Fig. 13b), low stomatal conductance (Fig. 13c), and a lower photosynthetic rate (Fig. 13d) were associated with decreased Ψ_50_, indicating a higher ability to withstand drought. The trade-off between xylem water permeability and vulnerability mediated by xylem anatomy has been well documented across Cupressaceae species, although it might not be a global pattern (Meinzer et al., 2009; Gleason et al., 2016). Among these axes, basal species (e.g., *Glyptostrobus*, *Taxodium*, *Metasequoia*) that grow in moist environments have lower Ψ_50_ with low wood density (Fig. 13a), higher xylem-specific conductivity (Fig. 13b), and higher area-based stomatal conductance and photosynthetic rate (Fig. 13c, 13d), whereas derived species growing in dry environments such as *Callitris*, *Juniperus*, and *Widdringtonia,* are on the opposite side of the axis (Fig. 13a,b,c,d).

**Fig. 13.**
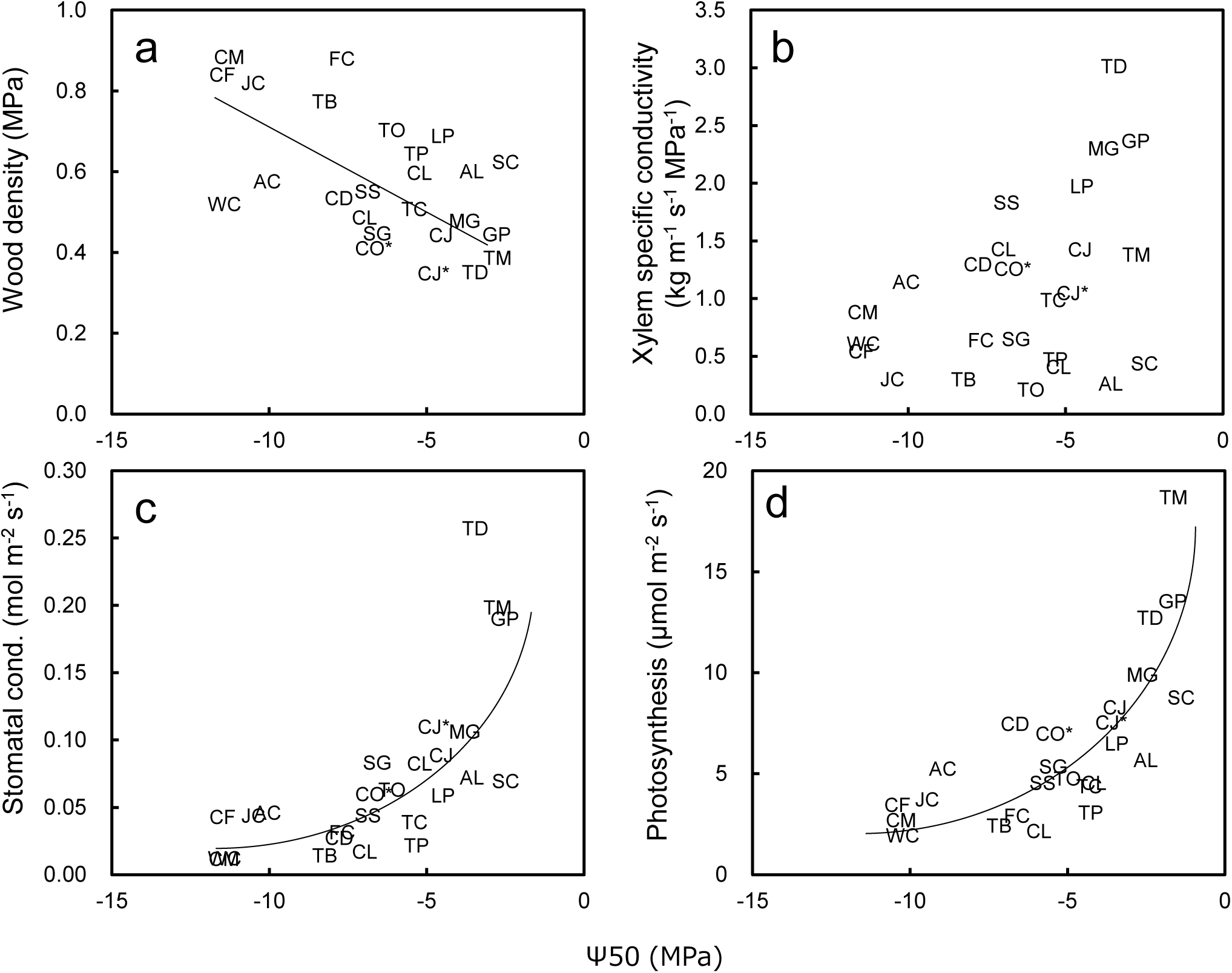
Relationship between leaf and stem hydraulic properties in Cupressaceae species. Abbreviations of species names are shown in Table 2. Species in bold type with an asterisk are from this study.

**Table 2.** Species name and abbreviations used in Figure 10. Data on *Cryptomeria japonica* and *Chamaecyparis obtusa* are from this study, and other data are from Pittermann et al. 2012.

| Species | Abbr. | Phenology, native habitat |
| --- | --- | --- |
| *Athrotaxis laxifolia* | AL | E, Montane forests, Tasmania |
| *Austrocedrus chilensis* | AC | E, The Andes, Chile and Argentina |
| *Callitris macleayana* | CM | E, Mesic-dry forests, eastern Australia |
| *Calocedrus decurrens* | CD | E, Montane forests, United States–Mexican Pacific Coast |
| *Chamaecyparis lawsoniana* | CL | E, Mixed forests, Oregon to northern California |
| *Chamaecyparis obtusa* (This study) | CO* | E, Mixed evergreen forests, Japan and Taiwan |
| *Cryptomeria japonica* | CJ | E, Mixed evergreen forests, Japan |
| *Cryptomeria japonica* (This study) | CJ* | E, Mixed evergreen forests, Japan |
| *Cunninghamia lanceolata* | CL | E, Mixed broad-leaved forests of southeast Asia |
| *Cupressus forbesii* | CF | E, Chaparral, southern California, northern Mexico |
| *Fitzroya cupressoides* | FC | E, Evergreen rainforest, Chile |
| *Glyptostrobus pensilis* | GP | D, Riparian, southern China |
| *Juniperus californica* | JC | E, Desert, southern California to northern Mexico |
| *Libocedrus plumosa* | LP | E, Mixed conifer rainforests, New Zealand |
| *Metasequoia glyptostroboides* | MG | D, Mesic mixed forests, central China |
| *Sciadopitys verticillata* | SC | E, Temperate moist forests, Japan |
| *Sequoia sempervirens* | SS | E, northern coastal California |
| *Sequoiadendron giganteum* | SG | E, Sierra Nevada, California |
| *Taiwania cryptomerioides* | TC | E, Cool temperate forests, Asia |
| *Taxodium distichum* | TD | D, Riparian regions in southeastern United States |
| *Taxodium mucronatum* | TM | D, Southern Texas, Mexico, Central America |
| *Taxus baccata* | TB | E, Broadly distributed across Europe |
| *Thuja plicata* | TP | E, Mixed coniferous forests, United States Pacific northwest |
| *Thujopsis dolabrata* | TO | E, Coastal and montane Japan |
| *Widdringtonia cedarbergensis* | WC | E, Fynbos, South Africa |

The traits of *C*. *japonica* and *C*. *obtusa* also fell on these correlation lines (Fig. 13). Among these axes, *C*. *japonica*, which is more basal than *C*. *obtusa*, was located next to the lowest Ψ_50_ (= the highest gas exchange) group, consisting of species of *Glyptostrobus*, *Taxodium*, and *Metasequoia*. This seems reasonable given that these genera form the same clade as *C. japonica*.

*C*. *obtusa*, which had a slightly lower Ψ_50_ than *C. japonica*, was located almost in the middle of the axes where close relatives, such as *Chamaecyparis lawsonia* (the same genus with *C*. *obtusa*) and *Thuja* and *Thujopsis* species, occurred. However, across all Cupressaceae species, the difference between *C*. *japonica* and *C*. *obtusa* was not large, nor was *C*. *obtusa* especially high in drought tolerance within the main Cupressaceae species. Japan, which is surrounded by the sea and has high-altitude mountains, has not suffered an extremely dry climate, even after the Cenozoic era, and only a few species of *Juniperus* with strong drought tolerance grow in alpine and coastal areas. Rather, there exist many relict genera, such as *Chamaecyparis, Thuja* and *Thujopsis,* which are located in the middle of the axis. This suggests that the mild climate of the Japanese archipelago became a refugia of these species with moderate drought tolerance in the arid Cenozoic (Farjon, 2008). However, high drought tolerance is generally achieved at the expense of a low growth rate. Therefore, the moderate drought tolerance of *C. obtusa* ensures a moderate growth rate and makes the species a major alternative to *C. japonica* at relatively dry forestry sites in Japan.

## 4 Implication for climate change

The detailed comparison of two major species highlights the importance of incorporating trait data into forest ecosystem models for more accurately predicting responses to climate change. Parameterizing models with various trait data enables us to obtain more realistic responses of trees and predictions. Recent studies demonstrate the limitations of plant functional type approaches and claim the need for trait-based approaches, for example, hydraulic responses (Anderegg, 2015). Most models at present are plant functional-type based; in other words, default parameters are prepared for different plant functional types; however, recent trait data compilation studies, such as this study, clearly demonstrate the diversity of traits among tree species even in the same plant functional type. The latest study even proposes flexible trait models for the next generation of vegetation models (Berzaghi et al., 2020). These trait-based approaches would succeed more easily in manmade pure forests than in mixed natural forests.

More specifically, our study provides important implications about possible differences in the species’ responses to climate change: no differences in SLA and photosynthetic ability per needle area were found between *C*. *japonica* and *C*. *obtusa*; on the other hand, the different structures of shoots and canopies between the two species may cause a difference in the amount of photosynthetic production per day. In particular, in the early planting period from just after plantation to canopy closure, *C*. *japonica m*ay have more advantages than *C*. *obtusa* due to its high photosynthetic production under bright light conditions. However, a big-leaf model, which is a typical photosynthesis model, does not explicitly describe the photosynthetic ability per part, structure or mass (Thérézien et al., 2007). For instance, values are the same for *C*. *japonica* and *C*. *obtusa* in the Biome-BGC model for most of the parameters. The response of stomata to climate change, such as the response of stomatal conductance to vapor pressure deficit (VPD), may differ between *C*. *japonica* and *C*. *obtusa*, but it is not clear at this stage due to a lack of data. In other words, it is possible to accurately grasp the type and amount of data that are lacking for modelling by using the database, which allows us to make an experimental plan to efficiently collect data according to a specific purpose. Regarding water use, we found various differences in related traits between *C*. *japonica* and *C*. *obtusa*; however, in general, most traits related to drought tolerance and xylem hydraulic safety except for gas exchange traits are rarely incorporated into models. This is perhaps partly because of the lack of data; drought tolerance and xylem hydraulic safety are new fields that have recently emerged.

In Japan, *C*. *obtusa* is a species planted in dry sites, while *C*. *japonica* is planted in wet sites; however, our compilation revealed that the drought tolerance of *C*. *obtusa* was moderate within the wide range of traits of the global Cupressaceae family. In fact, there are several reports of drought damage in *C*. *obtusa* forests, particularly in western Japan, which experiences more severe drought more often than in eastern Japan (Ogawa et al., 1996; Sanui et al., 1998). These facts imply that it is very likely that not only *C*. *japonica* but also *C*. *obtusa* will suffer from possible more severe droughts induced by future climate change.

## 5. Conclusion

The present study challenged the empirical knowledge of two contrasting plantation species in Japan. The intensive analysis of plant trait database clearly demonstrates that tree ecological traits recorded support traditional knowledge and empirical plantation management, such as preferable planting sites for both *C*. *japonica* and *C*. *obtusa*. Overall, even if the photosynthesis per foliage area is the same for *C*. *japonica*, photosynthetic production is higher due to the high shoot-level light utilization efficiency. In addition, high biomass allocation to the foliage and the low wood density of *C*. *japonica* result in a high stem volume yield. On the other hand, *C*. *obtusa* has high drought tolerance due to its lower transpiration rate, stomatal conductance and water potential at the foliar turgor loss point, whereas its photosynthesis at the shoot level is lower than that of *C*. *japonica*. These characteristics are consistent with traditional knowledge of suitable planting sites for both species (*C*. *japonica* on wet lower slopes and *C*. *obtusa* on dry ridges). Our finding that the most functional characteristics change according to tree age and/or height indicates that forest management also reflects a functional shift with the ontogeny of the tree. For example, to maximize tree production, fertilization to prevent foliar functional deterioration and thinning to compensate for the water supply can be considered with forest ageing. Furthermore, those approaches may also help to solve problems that cannot be addressed by empirical knowledge alone, i.e., the trait database is a powerful tool for modelling and predicting how forests respond to future temperature and precipitation associated with climate change, long-term cutting operations (e.g., more than 100 years), and more intense thinning to form mixed forests that contain broadleaf species. On the other hand, within the global Cupressaceae family, both species have moderate drought tolerance and photosynthetic rates, and the traits may be consistent with the historical climate of the Japanese archipelago being warm and humid without severe drought. The relatively low tolerance of both species may indicate a weak ability to withstand severe dry events associated with future climate change. This study clearly demonstrated that the plant trait database provides us a promising opportunity to verify out empirical knowledge of plantation strategies and help us to seek more climate change tolerant forest managements.

## 6 Author contributions

YO, SH and TK collected the data, analyzed the data, and drafted and edited the paper.

## 8 Acknowledgements

This study was funded as part of the project “Research on evaluation of influence of climate change on plantation in Japan” from the Ministry of Agriculture, Forestry and Fisheries.

